# Low-dimensional neural geometry underlies distributed goal representations in working memory

**DOI:** 10.1101/2023.12.07.570530

**Authors:** Mengya Zhang, Qing Yu

**Author notes:** **Correspondence should be addressed to: Qing Yu**, Institute of Neuroscience, Center for Excellence in Brain Science and Intelligence Technology, Chinese Academy of Sciences, Shanghai, 200031, China.

## Abstract

Successful goal-directed behavior requires the maintenance and implementation of abstract task goals on concrete stimulus information in working memory. Previous working memory research has revealed distributed neural representations of task information across cortex. However, how the distributed task representations emerge and communicate with stimulus-specific information to implement flexible goal-directed computations is still unclear. Here, leveraging EEG and fMRI along with state space analyses, we provided converging evidence in support of a low-dimensional neural geometry of goal information consistent with a designed task space, which first emerged in frontal cortex during goal maintenance and then transferred to posterior sensory cortex through frontomedial-to-posterior theta coherence for implementation on stimulus-specific representations. Importantly, the fidelity of the goal geometry predicted memory performance. Collectively, our findings suggest that abstract goals in working memory are represented in an organized, low-dimensional neural geometry for communications from frontal to posterior cortex to enable computations necessary for goal-directed behaviors.

## Introduction

In shifting environments, humans are capable of flexible cognition that relies on working memory, the ability to temporarily store and manipulate various kinds of information in mind [1]. These are not limited to basic sensory modalities but arguably more often involve abstract contents such as contingencies and contexts that guide goal-directed behaviors. When cooking without a recipe, one not only has to remember all the necessary ingredients (e.g., onions, celeries, and tomato, etc, for making a pasta), but the desirable state of each item (vegetables chopped, tomato crushed, and pasta cooked) and the action plans to achieve them (washing - chopping on a board - combining in a pan). All need to happen in a coordinated fashion. Yet within working memory research, the mechanisms of how neural codes of abstract task information and specific stimulus contents collectively support goal-directed behavior remains an open question.

The coding schemes of low-level sensory information in working memory have been relatively well-understood. Using multivariate decoding techniques or encoding models, stimulus-specific neural representations during memory delay has been found most strongly in sensory cortices responsible for the initial processing of corresponding sensory features [2–9]. These results highlight sensory cortex as a crucial site for stimulus maintenance in working memory [10]. In parallel, mounting evidence has suggested a prominent role of frontal cortex in representing task-related variables. Neural activity in frontal cortex has been found to preferentially encode higher-order representations of task contingencies [11], contexts [12], and rules [13–15], providing top-down control signals that target stimulus representations in downstream brain regions [10]. Despite the general dichotomy, abstract task representations are not constrained to frontal cortex, but are also observed in posterior sensory cortex [14–16]. These distributed task representations are enhanced during goal implementation compared to pure goal maintenance, which are accompanied by increased long-range functional connectivity [17] and may reflect a spreading of task information from frontal cortex during the active execution of control processes. Nevertheless, how the distributed task representations emerge and communicate with specific sensory information is still unclear.

It has been proposed that, to ensure stability in neuronal readouts and to enable task generalization and learning, population-level representations of task information may be compressed into a low-dimensional neural subspace [18,19]. Different cortical areas can interact through a selective low-dimensional communication subspace, which may potentially serve as a population-level neural mechanism for information relay between cortical regions [20,21]. Moreover, there is also evidence that low-dimensional control representations can guide the flow of information across brain areas without directing high-dimensional, detailed information [22,23], possibly through low-frequency, theta-band oscillations for long-range communications [17,24]. The findings summarized above suggest a potential mechanism by which abstract task information is organized in a low-dimensional representational format and relayed across cortical areas through subspace communication, possibly between frontal cortex and downstream sensory areas to support goal implementation.

In the present study, we set out to directly test this hypothesis by tracking the emergence of abstract task representations in different cortical regions and investigating how they interacted with specific stimulus contents during human working memory. Critically, we developed a principled approach to construct goal and stimulus representations in a structured manner, by designing a goal and a stimulus space, each defined by two orthogonal dimensions that formed a theoretical space. This design allowed for a direct examination of their neural representational geometries with respect to the theoretical ones, similar to how stimulus-specific representational subspaces were uncovered [25–28]. In two separate studies using electroencephalography (EEG) and functional magnetic resonance imaging (fMRI), healthy participants performed a novel delayed-recall task that required the maintenance of both a goal and a specific stimulus, and at a later stage the manipulation of the stimulus features based on the goal. Combining EEG with state space analyses, we found that task representations consistent with the low-dimensional goal space first appeared in frontal activity patterns before emerging posteriorly. Simultaneously the stimulus-related neural representations were present in posterior activity. The strength of the low-dimensional task and stimulus geometries was predictive of memory performance. Moreover, the transfer of goal representations from frontal to posterior regions and the interactions between goal and stimulus information were modulated by frontomedial theta-to-posterior coherence. Finally, using fMRI, the frontal low-dimensional goal geometry was localized to dorsolateral and medial prefrontal cortex (PFC), and the posterior counterpart, along with the corresponding stimulus geometry, was localized to visual regions. Together, these two studies provided converging evidence that low-dimensional, task-derived representational geometry of goal information emerges in frontal cortex and transfers to visual cortex, giving rise to successful goal-directed behavior.

## Results

### EEG behavioral results

Participants (n = 22) performed a working memory task on both memory goals and stimuli, alongside with EEG recording. Memory stimuli varied along two orthogonal feature dimensions (Figure 1B right), size (from small to big) and color (from green to red). Similarly, memory goals (Figure 1B left) also varied along two orthogonal goal dimensions, size (adjusting smaller to adjusting bigger) and color (adjusting greener to adjusting redder) goals. In other words, memory stimuli formed a two-dimensional (2-D) stimulus space, and memory goals formed a two-dimensional goal space. Prior to the main task, participants first learned the degree of required adjustment for all possible stimuli in the working memory task, such that for any given stimulus in the 2-D stimulus space, the correct degree of adjustment following a specific goal was pre-learned (Figure 1C). On each trial, participants were first cued with one of four possible goals (Goal cue: bigger and redder, bigger and greener, smaller and redder, smaller and greener; Figure 1A), and maintained the goal over the first delay period (Delay 1). After that they were presented with a to-be-memorized stimulus of a specific size and color (Sample), and maintained both the goal and stimulus over a second delay (Delay 2), before starting to make adjustments during the response period (Response; Figure 1E).

**Figure 1.**
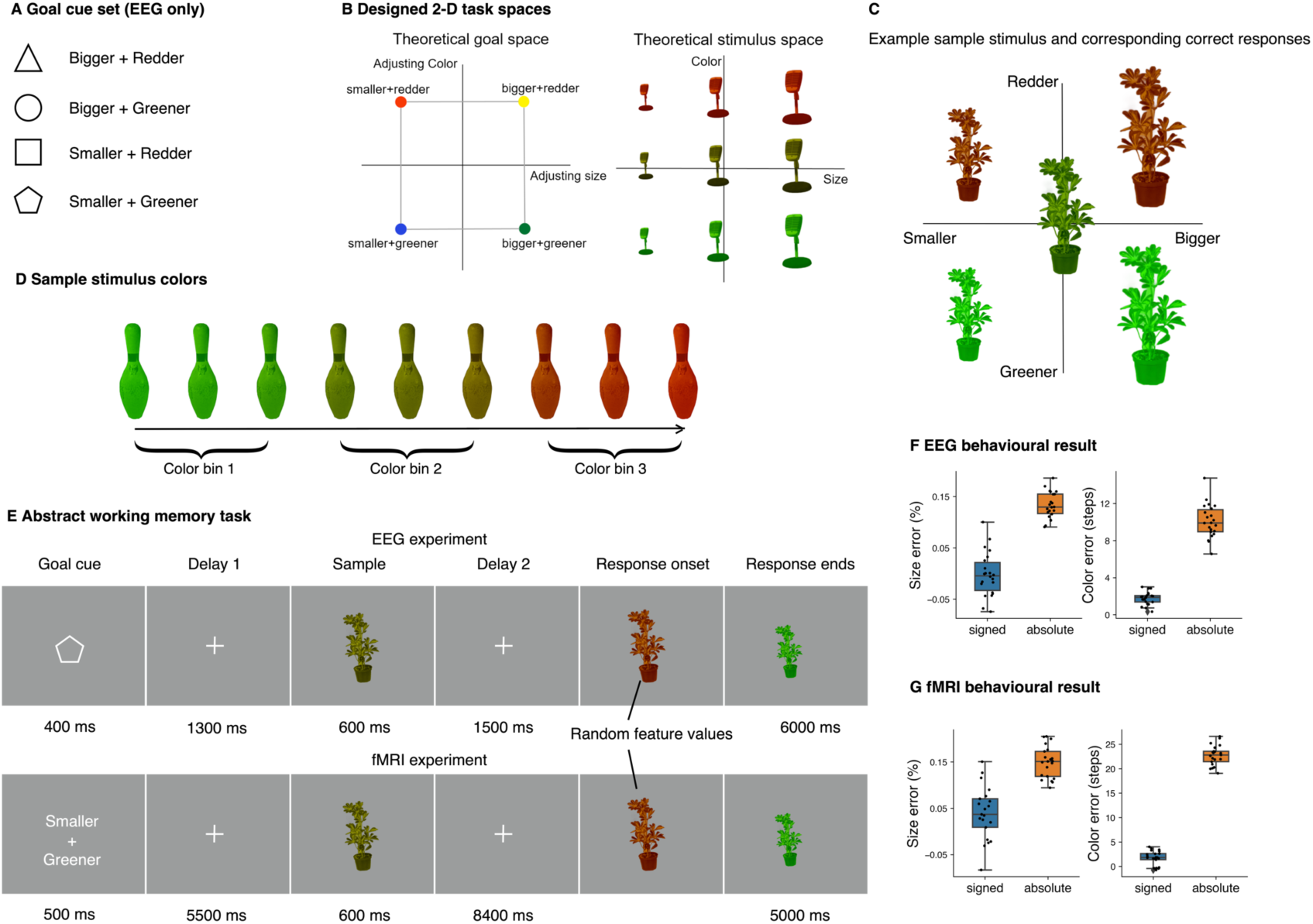
Task schematics and behavioral results. **(A)** Associations between shape cues and task goals. For EEG experiment, cues were used instead of text prompts for indicating task goals. Pairings did not change across participants. **(B)** Theoretical two-dimensional goal and stimulus spaces. To directly examine the neural geometries of working memory representations, both abstract and specific memoranda were constructed from two orthogonal axes. The task goal space consisted of adjusting size and color while the stimulus space of continuous stimulus size and color (red-greenness). Stimulus size and color were grouped into three bins each for subsequent analyses. **(C)** Example sample stimulus (center) and the corresponding correct answers (four quadrants) according to the task goals. The distances between any given sample and correct answers in terms of feature values were fixed and pre-learned by participants. **(D)** Illustration for all starting values in sample stimulus color. **(E)** Schematics of EEG and fMRI paradigms. The abstract working memory task required the memorization of task goal and stimulus features, and at response phase, the manipulation of stimulus based on the cued goal. The correct degree of manipulation was fixed and pre-learned by participants in a separate behavioral session. At the beginning of response, the appearance of a stimulus on screen marked the response onset, however, the size and color of this initial item were set with random values to prevent motor planning. **(F) & (G)** Size and color errors for EEG and fMRI experiments, respectively. The response errors were calculated by subtracting correct values from response. Mean signed error (blue) showed the degree of bias in participants’ responses whereas mean absolute error (orange) showed the degree of precision. Individual point represents mean error of each participant. Error bars denote 1.5 Inter quantile range (IQR).

Memory errors were calculated by subtracting correct answers from responses. Mean size error (Figure 1F) was −0.2% of starting size (SD = 4%), and was not significantly different from 0 (*t*(21) = 0.22, *p* = 0.82). Mean color error was 1.77 color steps (SD = 0.68), and was larger than 0 (*t*(21) = 12.17, *p* < 0.001). These suggested participants were able to perform the task according to the instructions and memorize the stimulus attributes, as there was no bias in size response and a small bias in color response towards redness, compared to the entire available color range (120 steps). We also showed the individual distributions of errors (Figure S1). Importantly, we used the absolute values of the response error in all subsequent behavioral correlation analyses, as the raw values were signed and only the magnitude itself measured performance. Mean absolute size error was 13.3% of starting size (SD = 2.6%), while mean absolute color error was 10.1 steps (SD = 1.81).

### Two-dimensional goal-specific representational geometry transfers from frontal to posterior channels

Having established behaviorally that participants could well maintain the remembered goals and stimuli, and manipulate the stimuli according to the remembered goal in the designed 2-D spaces, we next sought to explore the structures of goal and stimulus representations in the neural state space and their alignment with the designed 2-D task spaces. Recent work has successfully utilized dimension reduction techniques to reveal neural subspaces and representational geometries for stimulus information in working memory [26–29]. This approach transcends the mere confirmation of whether information is encoded (e.g., via multivariate decoding methods) and can provide richer descriptions of the coding principles adopted by neuronal populations in retaining and utilizing relevant information [30]. To this end, we used principal component analysis (PCA) to identify the two principal components (PCs) that explained the most variance in terms of task goals, in the group-concatenated data in each stage of a trial. We then examined whether the 2-D structural information in goals could be reflected in the 2-D neural subspaces as revealed by PCA. To examine representations of goal and stimulus, we primarily focused our subsequent analyses on two sets of EEG channels, frontal and posterior channels (Figure 2A). For visualization purposes, projected coordinates corresponding to unique goals were connected in the same order as in the designed goal space. To quantify the degree of which the representational geometry matched the 2-D structure, we computed a measure called circularity index (*C*), defined as the ratio between a geometric shape’s area and the square of the total length of perimeter. A larger circularity index would indicate higher similarity between the two structures. The result showed that frontal channel activities exhibited a neural geometric structure that followed the theoretical 2-D task space (i.e., adjusting size and color) during Delay 1 (*C* = 0.77, *p* = 0.02; Figure 2B). Meanwhile, although the hypothesized geometry (no intersected line) was also observed during other periods, none exceeded the critical statistical threshold derived from permutations (chance level circularity is approximately 0.74 with variations across epochs; Goal cue: *p* = 0.29; Sample: *p* = 0.19; Delay 2: *p* = 0.38; Figure S2A top). For posterior channels (Figure 2B bottom), activity patterns did not exhibit the representational structure predicted by the task space, at least not in the subspace defined by the first two PCs (Goal cue: *p* = 0.51; Delay 1: *p* = 0.94; Sample: *p* = 0.81; Delay 2: *p* = 0.85; Figure S2A bottom). However, when looking more closely at the changing dynamics of circularity index instead of averaging data over stages (see next paragraph), it could be seen that the strength of 2-D goal geometry increased towards the late Delay 2, which was indeed significant if these time points (3600-3800 ms) were considered (*C =* 0.75, *p* = 0.04).

**Figure 2.**
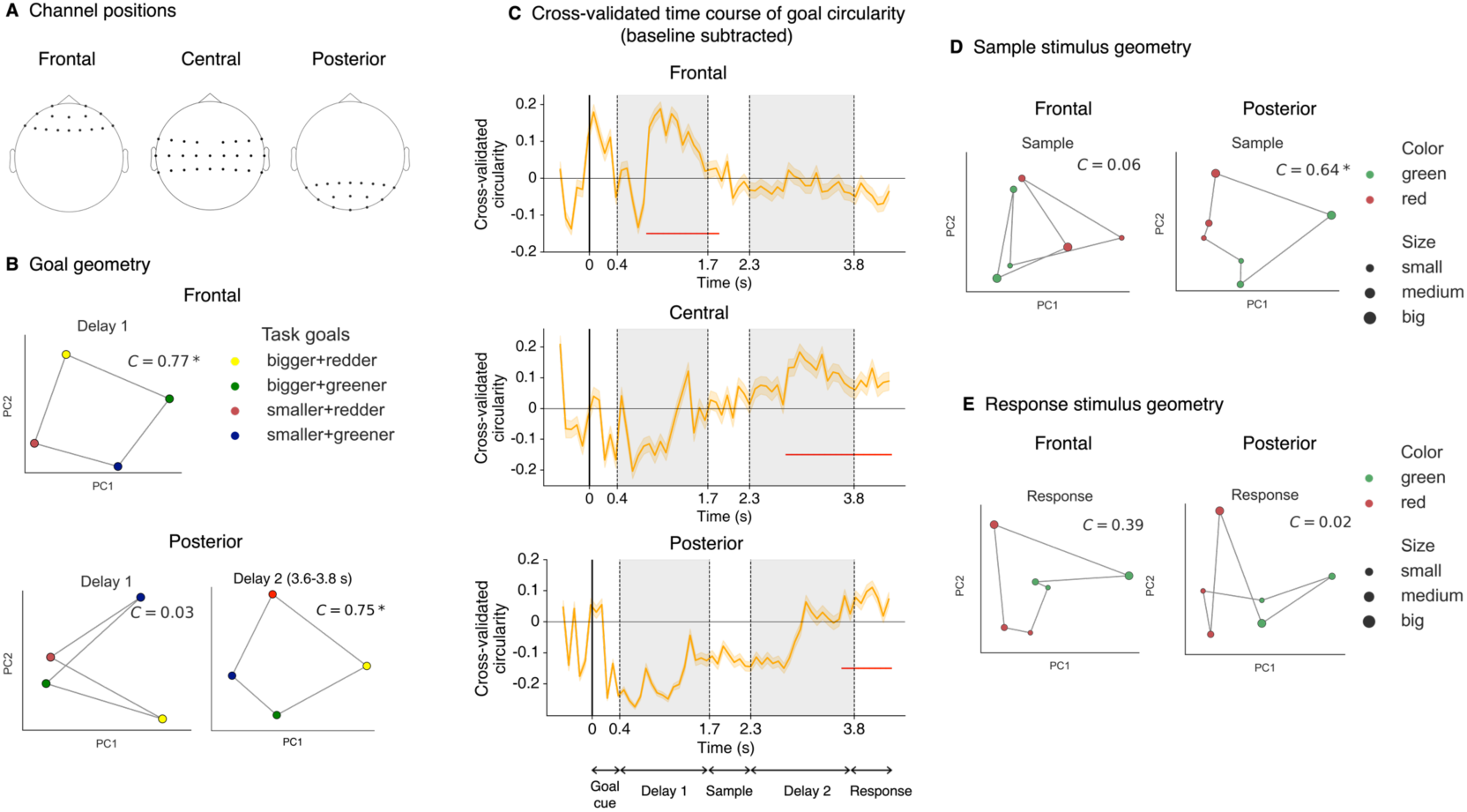
Two-dimensional representational geometries of goals and sample stimuli. **(A)** Demonstration of frontal, central and posterior channel locations. **(B)** Task goal conditions projected onto the 2-D neural subspace identified by group-level PCA from averaged Delay 1 period for frontal channels (top) and posterior channels for Delay 1 and late Delay 2 (bottom). *C* denotes circularity index. Asterisk denotes significance from permutation testing (*p* < 0.05). **(C)** Time courses of cross-validated and baseline-subtracted circularity index using data averaged over a non-overlapping 80-ms sliding window. Positive values denote significant circularity index (baseline-subtracted). Error bars denote 95% confidence interval derived from bootstrapping procedures. Red horizontal lines denote time points when 2-D goal geometry was significant. **(D)** Frontal and posterior stimulus-specific 2-D geometries for the Sample epoch. Stimulus feature values were aggregated into three bins and the middle color bin were discarded in order to calculate circularity index, resulting in six unique stimulus conditions. **(E)** Similar to **(D)** but for responded stimuli, which was calculated using the feature values after participant have finished adjusting (i.e., answers).

The hypothesized 2-D goal structure was present firstly at frontal channels during goal maintenance (Delay 1) then at posterior ones during goal implementation (Delay 2), which bear the question of whether the two neural geometries were related or independently formed. To provide insight into this, we included the rest of the channels that sit between frontal and posterior areas (i.e., central channels). By comparing the time-resolved circularity index, we could observe the temporal relationship formed by the changes in the strength of 2-D goal geometry across these groups of channels. Cross-validated time courses of circularity index (from bootstrapping trial-wise data in a stratified manner) using a non-overlapping 80-ms sliding window were computed. The hypothesized 2-D goal geometry was present in all three groups of channels albeit at different time periods over the course of a trial, with frontal activities exhibiting it first in Delay 1 (820-1780 ms; Figure 2C), followed by central channels which started showing significant time points as early as Delay 2 onset (2820-4260 ms); both temporally preceded posterior activities in which 2-D goal geometry only arose towards the end of Delay 2 (3620-4300 ms). Therefore, this suggests the 2-D goal geometry was first formed in frontal channels and emerged over time to posterior channels, possibly by travelling backwardly through central channels.

To rule out the possibility that motor preparations caused the observed 2-D representations, as a result of the four available button-feature adjustment mappings remaining the same across trials and participants, we correlated the goal and motor-specific circularity time courses (see Methods), which was not significant in either frontal or posterior activities (frontal: *r* = −0.08, *p* = 0.54; posterior: *r* = 0.11, *p* = 0.39), indicating that representations of motor responses could not have led to the 2-D goal geometry observed in the data. Moreover, the cross-validated circularities for motor representations were calculated and there was a brief period when frontal channels showed 2-D geometry (Figure S3). However, we reasoned that because none of the channels manifested any significant result towards Response, which was the critical period if motor preparation signals existed; the results here within Delay 1 was in fact a spillover from the 2-D goal representations, as motor response and task goals were indeed related in our post-hoc analysis (*χ*^2^ = 588.2, *p* < 0.001).

**Figure 3.**
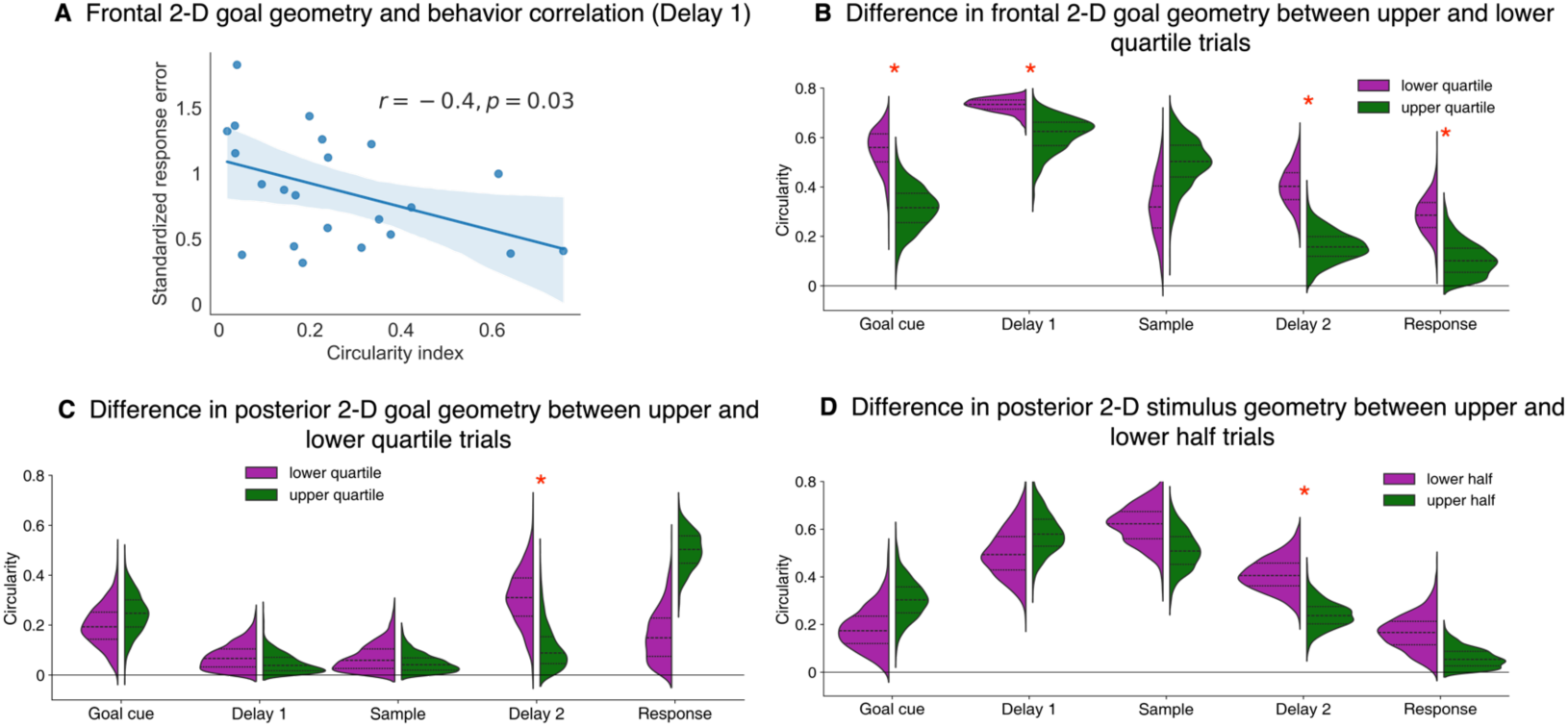
(A) Significant correlation between frontal 2-D goal geometry in Delay 1 and the averaged response error. **(B)** Difference in frontal 2-D goal geometry between trials in the upper and lower quartiles based on response errors. Lower (purple) and upper (green) quartile correspond to trials with smaller and larger errors, respectively. Red asterisk denote significance at α = 0.05. **(C)** Same as **(B)** but for posterior goal geometry. **(D)** Difference in posterior 2-D stimulus geometry using a median-split.

To verify that PCA results based on group-concatenated data still held true at individual level, and that the 2-D goal representational geometry measured by circularity index was also comparable with other more traditional metrics, we computed representational similarities between theoretical models and our EEG data. If the group-concatenated result truly indicated that individual goal representations followed the 2-D geometry, then it should be manifested in a subject-wise representational similarity analysis (RSA) using the corresponding 2-D goal model (Figure S4A). This is exactly what we saw: only during Delay 1 in frontal channels, and late Delay 2 onwards for posterior channels, time-resolved RSA correlations were significant, in line with the 2-D goal geometry revealed by group PCA, which was consistent even at the individual level. Thus, we confirmed that the results above were robust and commensurable with the alternative analytic approach of RSA.

### Strength of 2-D goal geometry predicts memory performance

Having established the existence of the 2-D goal-specific representations in both frontal and posterior channel activities which possibly cascaded from front to back over time, we tested whether the strength of such neural geometry was related to behavior. Firstly, correlational analyses were performed between individual circularity indices and memory performance as measured by the sum of standardized absolute color and size errors. We hypothesized that a stronger 2-D goal geometry would lead to a better separation of conditions for downstream populations and more robust spatial transmission of information, hence better performance should be observed. Consistent with our prediction, 2-D goal geometry in frontal channels was negatively correlated with response error during Delay 1 (Spearman’s correlation; *r* = −0.40, *p*= 0.03; one-sided test; Figure 3A), but not in posterior channels (*r* = 0.24, *p* = 0.85). The correlation analyses demonstrated that the degree to which individuals formed frontal low-dimensional goal representations was predictive of the overall performance.

We also performed within-subject comparisons which afford more sensitivity than between-subject methods as they are less subject to individual differences that may affect task performance. For goal geometry, we took the bottom and top quartiles of each participant’s trial-wise data based on the magnitude of combined response error to compute group-level circularity indices corresponding to the good and bad trials within each epoch, respectively. Specifically, the two sets of data were further bootstrapped in order to achieve a more stable estimation for the difference in 2-D geometry strength. In line with our prediction, trials with smaller errors were associated with stronger 2-D goal geometry than trials with larger errors. In frontal channels, this was true for Goal cue (*p* = 0.005), Delay 1 (*p* < 0.001), Delay 2 (*p* < 0.001) and Response (*p* = 0.01). For posterior activities, although we did not find a correlation with behavior, there was a significant difference in 2-D goal geometry for Delay 2 (*p* = 0.03), although the effect was reversed at Response (*p* = 0.001), showing a less consistent trend compared to frontal channels. Overall, these behavioral analyses support the functional relevance of the 2-D goal geometry in working memory.

### Posterior activities also exhibit 2-D stimulus-specific geometry

In addition to the representations for task goals, what is the geometric structure for stimuli which themselves were drawn from a feature space consisting of two independent dimensions? Based on previous work using simple visual features that demonstrated low dimensionalities [26–29], it is expected that similar to goal representations, a low-dimensional subspace can well capture the stimulus-dependent variances in neural activity. When applying PCA to identify such subspace specific to stimulus color and size, we observed the 2-D stimulus geometry in posterior channels. The structure followed theoretical geometry (borders connecting neighboring coordinates did not cross each other) during sample presentation (*p* = 0.007; Figure 2D) and unexpectedly during Delay 1 as well (*p* = 0.009), but not at other periods (Goal cue: *p* = 0.16; Delay 2: *p* = 0.09; Figure S2B). We argue that the result in Delay 1 cannot be explained by a correlation between goal and stimulus, as they were fully counterbalanced by design (*χ*^2^ = 0.03, *p* > 0.99). Moreover, our subsequent analysis further demonstrated that the within-subject-level circularity in Delay 1 failed to predict behavior (see next paragraph), indicating that the result in Delay 1 was likely due to sporadic fluctuations. In contrast, in frontal channels 2-D stimulus structure was not observed throughout the trial (Goal cue: *p* = 0.51; Delay 1: *p* = 0.79; Sample: *p* = 0.77; Delay 2: *p* = 0.51; Figure S2B), suggesting that stimulus-specific processing and maintenance were executed mainly by regions monitored by posterior channels. Of note, stimulus size and color were grouped into three bins each, resulting in nine unique stimulus-specific conditions. In the current result we used six of them in order to construct a close-formed geometric shape to calculate the circularity index (i.e., the middle row in Figure 1B right was discarded).

Given that a 2-D stimulus geometry was identified in posterior activities, it stands to examine whether such representations were related to behavior. Although no group-level correlation was found at any epoch, there was a difference in stimulus-specific circularity index using trials median-split into larger and smaller response errors. This effect was significant for Delay 2 (*p* = 0.003; Figure 3D), but not for the rest of the epochs (Goal cue: *p* = 0.92; Delay 1: *p* = 0.79; Sample: *p* = 0.09; Response: *p* = 0.08), further illustrating that both 2-D stimulus and goal representations contributed to the final behavioral performance, although the latter exhibited a more robust influence.

In addition, to successfully adjust the sample features to the correct degree, participants might have preemptively manipulated the remembered stimulus in mind in preparation for the responses from Delay 2 onwards. To test this hypothesis, we performed the same PCA procedure but binned the trials based on the responded feature values, and found that there was a trend for 2-D structure of the responded stimulus in frontal activities at the time of response (*p* = 0.07) but not for other epochs (Sample: *p* = 0.15; Delay 2: *p* = 0.14; Figure 2E and S2D). No 2-D structure was found in posterior activities at any time (Sample: *p* = 0.64; Delay 2: *p* = 0.61; Response: *p* = 0.92).

### Long-range theta connectivity mediates the backward transmission of 2-D goal geometry and integration with stimulus information

So far, we have observed a distributed network for goal representations in which goal geometry transferred front to back in a low-dimensional format, and on top of that, low-dimensional stimulus representations in posterior channels. However, what mechanisms mediate the backward transfer of 2-D goal geometry and whether this process was related to the integration of goal and stimulus information in working memory? To this end, we targeted frontomedial theta (FMT) to posterior coherence as a measure of time-resolved long-range connectivity [31], and examined whether FMT was predictive of the strength of neural geometries for task goals, stimulus and responses. Firstly, two significant frontomedial-to-posterior coherence clusters (see Methods) were identified that covered the delta and theta frequencies (Figure 4A), one spanning from 0 – 1276 ms (Goal Cue and Delay 1), the other from 1728 – 3268 ms (Sample and Delay 2). As the next step, we used the theta range (4-7 Hz) of these two group-level clusters as masks to extract individual FMT strength and examined its relationship with task representations and behaviors.

**Figure 4.**
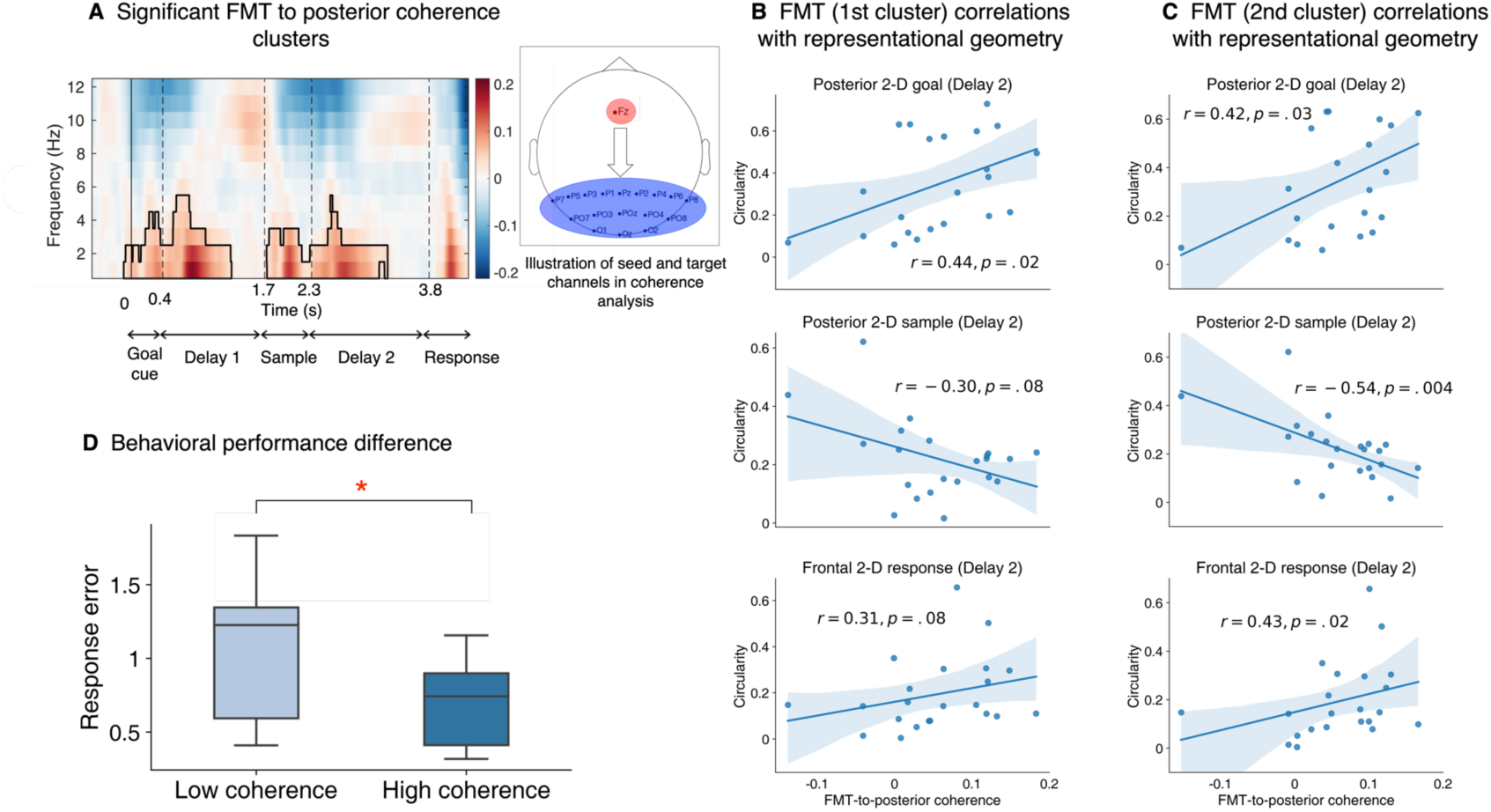
FMT to posterior coherence and correlations with representational geometries. **(A)** Time-resolved frontomedial (channel Fz)-to-posterior coherence between 1-12 Hz. Solid black line encircles the significant time-frequency clusters. For behavioral correlations, only sub-clusters falling within the 4-7 Hz (theta band) were used as masks to extract individual coherence strength. **(B)** Correlation results (Spearman’s *r*) between the first FMT coherence cluster and representational geometries (i.e., circularity index) of task goals (upper), sample stimulus (middle) and response (lower) at Delay 2. Each dot represents an individual. Note that the spatial locations of significant correlations varied across the types of representation. **(C)** Same as **(B)** but using the second FMT coherence cluster. Correlation patterns were consistent between the two clusters, suggesting similar function of the FMT coherence at both times. **(D)** Difference in behavioral performance between individuals with low and high FMT coherence strengths in the second cluster.

The first FMT cluster (Figure 4B 1^st^ row) was predictive of posterior 2-D goal geometry at Sample (Spearman correlation, one-tailed; *r* = 0.51, *p* = 0.007) and Delay 2 (*r* = 0.44, *p* = 0.02); there was no evidence for such a relationship during Goal cue (*r* = −0.09, *p* = 0.66) or Delay 1 (*r* = −0.008, *p* = 0.51). This suggested FMT to posterior connectivity could underlie the transfer of 2-D goal representations to posterior sites which initially emerged frontally and provided a tentative explanation for the temporal cascading relationship observed earlier across frontal, central and posterior channels. Moreover, the same effect was also found for the second FMT cluster (Figure 4C 1^st^ row; Goal cue: *r* = −0.32, *p* = 0.92; Delay 1: *r* = −0.11, *p* = 0.69; Sample: *r* = 0.45, *p* = 0.02; Delay 2: *r* = 0.42, *p* = 0.03), which could indicate both FMT clusters share the function of relaying the 2-D goal information backwards.

We next wondered whether FMT was also related to the neural geometries of sample and response stimulus representations, as one potential function of long-range fronto-to posterior connectivity is the transfer of task-related control signal to modulate stimulus processing. Based on this and our finding that FMT is involved in goal information relay, we hypothesized that FMT was related to the integration of goal and sample information, and therefore the formation of response representations as the product. Specifically, FMT should be negatively correlated with sample, and positively corelated with response geometries, given that stronger goal representations should be associated with the rise of response representation and the decay of sample representations (due to transformation of original stimulus information). Aligned with the predictions, the second FMT cluster was negatively correlated with the 2-D sample structure posteriorly at Delay 2 (Figure 4C 2^nd^ row; *r* = −0.54, *p* = 0.004), although its correlation with posterior response structure was not significant (*r* = 0.02, *p* = 0.45). Nevertheless, recall that we observed marginal evidence for response representations in frontal instead of posterior channels, we thus examined the relationship between the second FMT cluster and response structure in frontal channels, and found the two were indeed positively correlated (Figure 4C 3^rd^ row; *r* = 0.43, *p* = 0.02) at Delay 2. As such, the localization of the relationship was coherent with that from the PCA subspace analyses where posterior sample representations and frontal response representations were identified. Moreover, similar but weaker patterns were observed between the first FMT cluster and the sample and response geometries (Figure 4B 2^nd^ row; *r* = −0.30, *p* = 0.08; Figure 4B 3^rd^ row; *r* = 0.31, *p* = 0.08, respectively). The temporal discrepancy (first FMT cluster took place in Goal cue and Delay 1) could indicate that stimulus transformation also depended on the initial strength of goal information relay.

Given the importance of transferring goal representations to influence stimulus processing in the present task, we investigated the relation between FMT connectivity and behavioral performance. Participants were median-split based on the FMT coherence strength and between-group difference in response error was tested. Those with stronger connectivity during Sample and Delay 2 (i.e., the second cluster) performed better overall, *t*(20) = 2.17, *p* = 0.02 (Figure 4D). On the contrary, there was no difference if participants were divided based on the first FMT cluster, *t*(20) = 0.79, *p* = 0.22. This distinction could be a result of the timing of the second FMT cluster being closer to the arrival of stimulus information and the implementation of task goals, hence more linked with the underlying computations than the first one. Collectively, these findings illustrate that long-range connectivity is involved in the relay of goal information in a low-dimensional format, which is in turn critical to the integration with the veridical stimulus to produce final responses.

### FMRI behavioral results

To further localize brain regions with the designed goal and stimulus geometries, we conducted an fMRI experiment using the same task (n = 21), with event timing adjusted to compensate for the sluggishness of BOLD signals. We first assessed participants’ behavioral performance in the fMRI experiment following the behavioral measures in the EEG experiment. Overall, results from the fMRI experiment were comparable to those from the EEG experiment: mean size error was 4% of starting sample size (SD = 5%), and significantly different from 0 (*t* (20) = 3.20, *p* = 0.004). Mean color error was 1.76 steps (SD = 1.45), and larger than 0 (*t* (20) = 5.59, *p* < 0.001). Mean absolute size error was 14.9% of starting size (SD = 3.3%), while mean absolute color error was 22.7 steps (SD = 2.03).

### Two-dimensional goal geometry in dorsolateral prefrontal, medial prefrontal and visual cortex

In the EEG study, frontal and posterior channels displayed differential sensitivity to goal and stimulus representations. These signals likely arose from frontal and visual cortex, respectively. To have a finer understanding of the spatial localization of the 2-D representational geometry associated with goal and stimulus space, we divided PFC and visual cortex into regions-of-interest (ROIs) based on the Human Connectome Project (HCP) atlas [32], and examined the representational structures of goals and stimuli in these ROIs (Figure 5A and 5E), using trial-wise beta coefficients estimated for each task epoch of interest from general linear models (GLMs).

**Figure 5.**
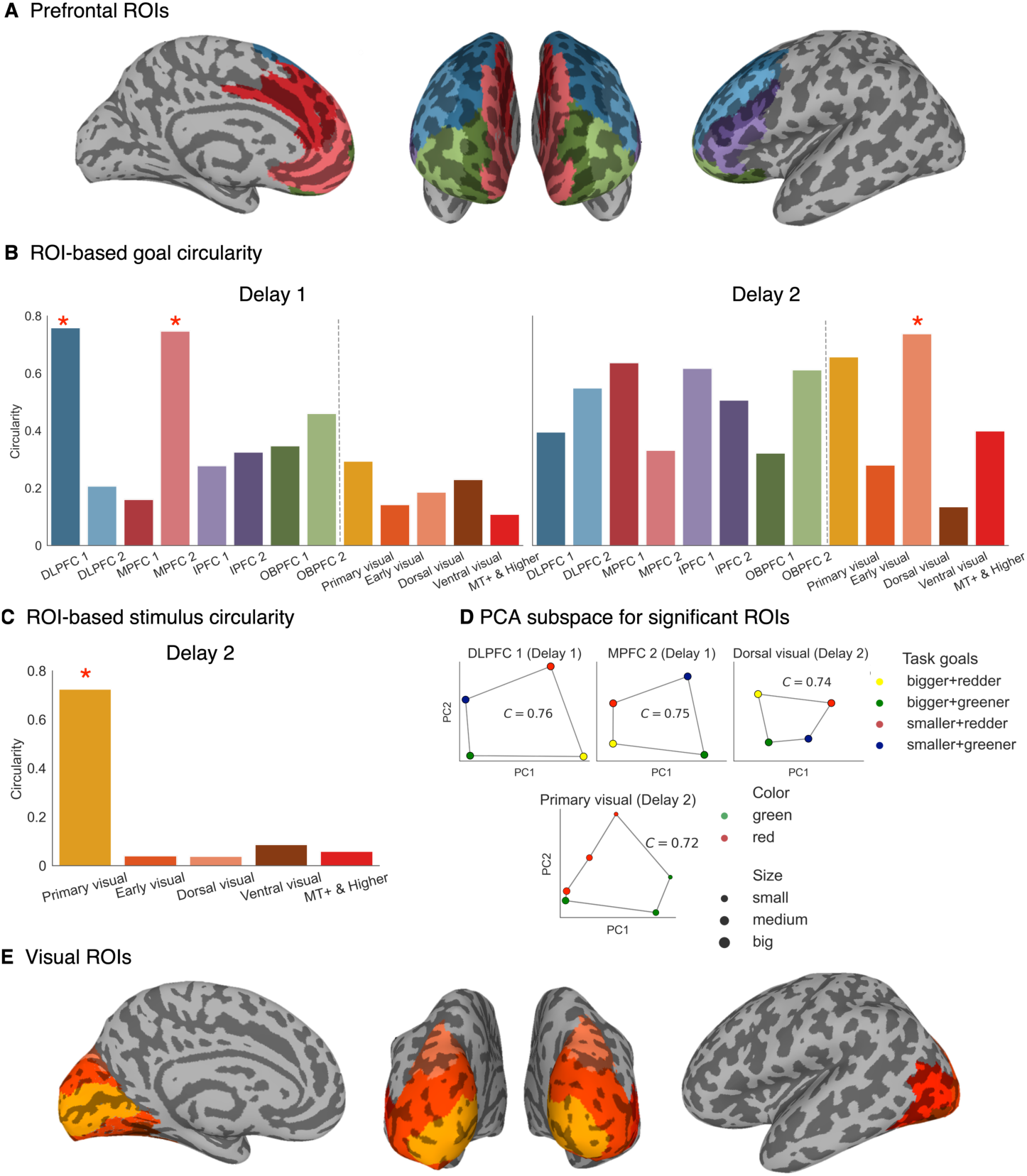
FMRI ROI-based neural geometry results. (A) PFC parcellations. Definition for ROIs was based on the HCP atlas. Each section was divided into two ROIs: dorsolateral PFC (DLPFC 1 and 2), medial PFC (MPFC 1 and 2), inferior PFC (IPFC 1 and 2) and orbitofrontal PFC (OBPFC 1 and 2). (B) Circularity indices for task goals using beta coefficients of all voxels within an ROI estimated from trial-wise GLMs. Red asterisk denotes *p* < 0.05 based on permutations. (C) Same as (B) but for stimulus circularities in visual ROIs at Delay 2. (D) The subspace plots depict the representational structures of ROIs with significant results for goals (top) and samples (bottom). (E) Visual cortex parcellations. These ROIs include Primary visual cortex, Early visual/extrastriate cortex, Dorsal visual cortex, Ventral visual cortex and Middle temporal complex (MT+) and higher visual areas.

We found consistent evidence for the 2-D goal structure in frontal regions during Delay 1 and in visual regions during Delay 2, confirming the results from our EEG experiment. In Delay 1, neural activities in a dorsolateral PFC (DLPFC) cluster (lateral superior frontal gyrus; DLPFC 1) and a medial PFC (MPFC) cluster (medial section of superior frontal and orbital section of middle frontal gyri, and part of anterior cingulum; MPFC 2) showed the 2-D structure predicted by the goal space (Figure 5B left and 5D top; DLPFC 1: *p* = 0.03; MPFC 2: *p* = 0.04). Oppositely, visual areas, and specifically the dorsal visual areas (superior and middle occipital gyri) exhibited this geometry at Delay 2 (Figure 5B right; *p* = 0.049). The alternative RSA approach was again performed to compare the neural data with hypothesized 2-D model of task goals on an individual level. Corroborating the PCA-based result, both frontal ROIs were related to the 2-D model at Delay 1 (DLPFC 1: *t*(20) = 2.72, *p* = 0.01; MPFC 2: *t*(20) = 2.30, *p* = 0.03; Figure S4B); however, we did not replicate the finding in the dorsal visual ROI at Delay 2 (*t*(20) = 0.73, *p* = 0.47). This inconsistency could be due to that the RSA approach assesses representational similarity in a higher-dimensional space beyond the first two PCs. As for the 2-D stimulus-specific geometry predicted by color and size features, it was identified in the primary visual cortex among all posterior ROIs (Figure 5C and 5D bottom; *p* = 0.009). To ensure that the results was not affected by the number of voxels used in ROIs, we selected the 500 most positively activated voxels for Delay 1 and 2 respectively within each ROI and repeated the above analysis. The results remained largely unchanged (Figure S5). Taken together, data from the two experiments demonstrate the spatial and temporal characteristics of the 2-D geometries are in fact robust and can be uncovered by different recording modalities.

Lastly, given that the EEG experiment showed a trend for 2-D response representation in frontal activities which seemed to be modulated by FMT coherence, we also sought to clarify this with fMRI. The same ROI-based approach was taken to calculate the response-specific circularities. The middle frontal gyrus (DLPFC 2; *p* = 0.03) and early visual/extrastriate cortex (*p* = 0.03) exhibited the 2-D response geometry (Figure S6) in the late Delay 2, that is, the second half of the period. However, only visual areas including the primary visual, early visual and to a lesser degree middle temporal visual areas held this low-dimensional format of transformed stimulus representations during Response (primary visual: *p* = 0.003; early visual: *p* = 0.003; MT+ and higher visual: *p* = 0.04). Overall, both frontal and visual ROIs demonstrated the 2-D response geometry in the fMRI experiment.

**Figure 6.**
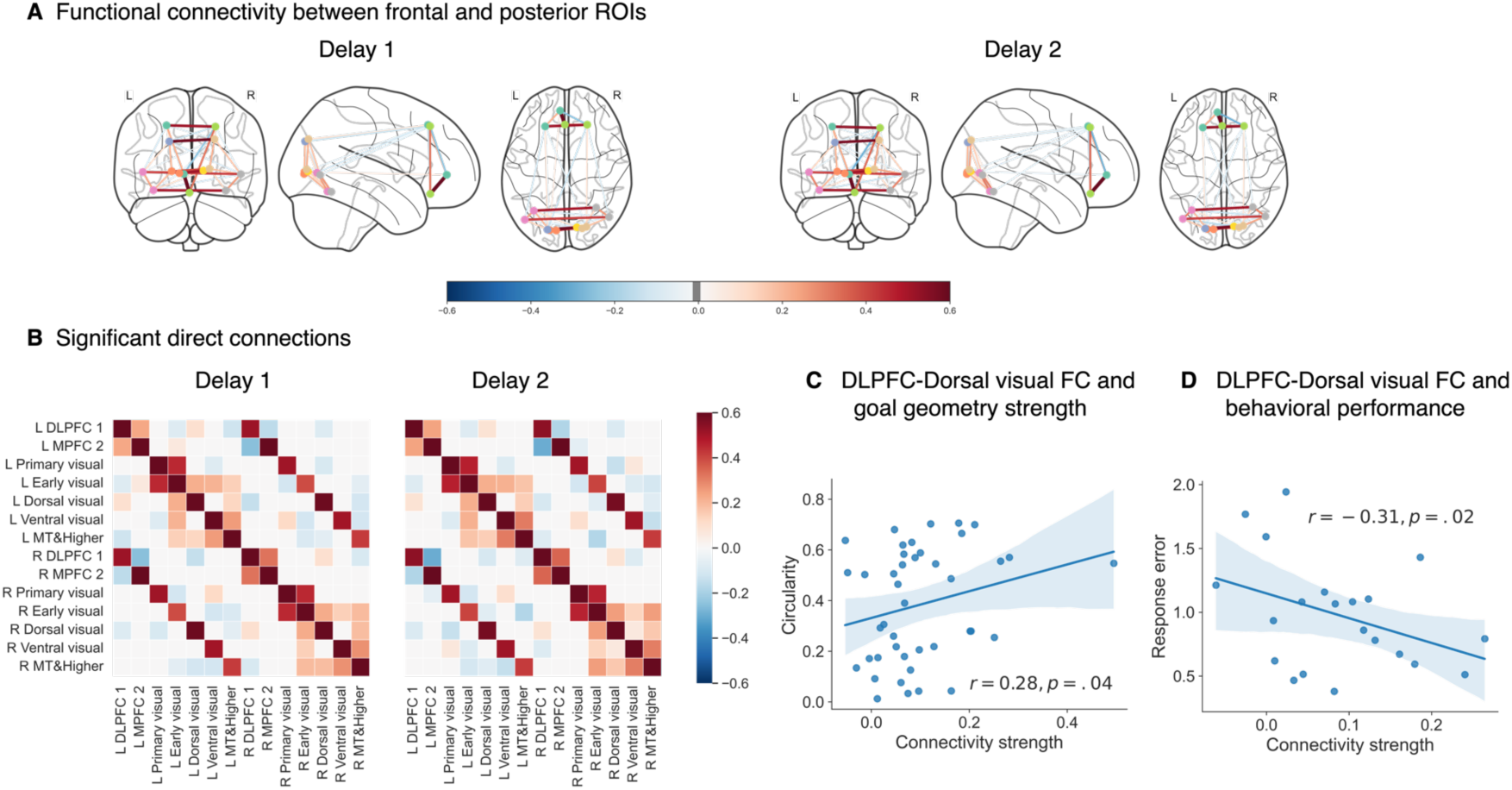
Functional connectivity among selected ROIs. **(A)** Significant connections (partial correlations) for Delay 1 and Delay 2 using ROIs showing significant frontal goal geometries (DLPFC 1 and MPFC 2) and all posterior visual ROIs. Red denotes positive correlations and blue denotes negative correlations. **(B)** Matrices showing significant pairwise connections. Of note, cells without colors did not pass the significance test corrected for multiple comparisons. Left and right side of the same ROI as well as areas of spatial vicinity tended to be positively correlated. **(C)** The connection strength between DLPFC 1 and dorsal visual cortex in Delay 1 was predictive of goal circularities within the latter in Delay 2. Both ROIs showed significant 2-D goal geometries in Delay 1 and 2, respectively. **(D)** Higher connection strength between DLPFC 1 and dorsal visual cortex in Delay 2 was predictive of better behavioral performance.

### Functional connectivity between DLPFC and posterior regions mediates 2-D goal geometry and predicts task performance

Having identified the regions exhibiting the 2-D structures related to task goals and sample stimuli, we evaluated whether long-range connectivity between these particular regions mediated the transfer of goal representations from frontal to posterior areas, similar to what was observed in the EEG experiment. We used the trial-wise beta estimates of delay-period activities to compute partial correlations among pairs of regions to assess direct connections, controlling for influence from other sources [33,34]. For this analysis, we parcellated the bilateral ROIs into unilateral ones to take into account the variations in ipsi- and contra-lateral connections (Figure 6A). The prefrontal ROIs (DLPFC 1 and MPFC 2), which exhibited significant 2-D goal geometry in Delay 1, generally showed significant positive correlations among themselves and a robust bilateral positive relationship with the dorsal visual areas, which showed 2-D goal geometry in Delay 2 (Figure 6B). Otherwise, long-range fronto-occipital connections were sparse and negative if existed.

In light of the EEG connectivity result that long-range connectivity mediated the relay of goal information, a similar approach in the fMRI domain was employed. The advantage of fMRI allowed us to further limit this test to ROIs specifically holding the 2-D goal geometry, namely DLPFC 1, MPFC 2, and dorsal visual areas. We only used ipsilateral connections, for instance, between left/right DLPFC and left/right dorsal visual areas, given there is evidence supporting both structural and functional connectivity on the same side [35,36]. The direct connection between DLPFC 1 and dorsal visual areas at Delay 1 was predictive of 2-D goal geometry in the latter ROI at Delay 2 (Figure 6C; Spearman correlation; *r* = 0.28, *p* = 0.04). The timing of the relationship was partially in accordance with our EEG finding, suggesting a delayed usage of goal representation after stimulus encoding. We did not find the same relation using the MPFC 2 ROI as the frontal node (*r* = −0.05, *p* = 0.63), despite the fact that it also held a copy of the 2-D goal representations at Delay 1. Additionally, consistent with the EEG result, connectivity strength between DLPFC 1 and dorsal visual areas at Delay 2 was also predictive of task performance (*r* = −0.31, *p* = 0.02; Figure 6D). In short, the fMRI connectivity analyses replicated the findings of the EEG experiment, that is, the long-range connectivity was associated with backward transmission of goal information in the 2-D format which subsequently influenced behavior.

## Discussion

In the present study, we investigated the distributed goal representations in working memory, using a newly-developed behavioral paradigm that required retention of both task goals and stimulus contents in designed two-dimensional goal and stimulus spaces. Leveraging recently advanced state space analyses, our first EEG experiment revealed that the representational geometry of goals followed the theoretical two-dimensional structure. This two-dimensional goal geometry first emerged in frontal channels during goal maintenance and then transferred to posterior channels for goal implementation. Notably, the fidelity of the goal geometry was predictive of individual memory performance. Meanwhile, a two-dimensional stimulus geometry was observed in posterior channels in accordance with the theoretical stimulus structure. The frontal goal geometry transferred to posterior sites and interacted with the posterior stimulus geometries through FMT coherence. A second fMRI experiment further indicated that both DLPFC and MPFC exhibited the desired two-dimensional goal geometry. However, only DLPFC demonstrated significant functional coupling with posterior visual cortex that predicted the transfer of goal structure and behavior. Collectively, our findings suggest a potential neural mechanism for how frontal and posterior cortices are orchestrated through communications of low-dimensional geometry to implement computations necessary for goal-directed behavior.

### Low-dimensional representations for transferring task information

Frontal cortex has long been considered as a core region for processing task-related variables, with aggregated BOLD activity that reflects different levels of abstraction during cognitive control [37,38], and decoding of a variety of abstract task information [14,15,39,40]. Beyond the focus of frontal areas, many studies have also reported successful decoding of task sets in targeted sensory cortex [14,16]. We directly addressed the roles of the distributed representations of abstract task information, and how they support goal-directed behavior in working memory by tracking the representational geometries of structured task information in the brain. During the first delay when goals were encoded but not yet implemented, task goals awaiting to be integrated with specific contents in working memory were indeed maintained in a compressed format of representations that reflected the designed dimensions. Importantly, this low-dimensional goal geometry was predictive of subsequent memory performance and cannot be accounted for by preparative motor planning signals. These together suggest that at the population level, frontal cortex represents abstract goal information in a manner similar to how sensory cortex encodes feature information [25,26,29], despite neurons in the two cortices possess distinct tuning properties (i.e., mixed vs. simple selectivity). In other words, frontal cortex organizes goals into an abstract relational space based on their underlying structure, which can be maintained in working memory for later control of stimulus adjustments [41]. This is reminiscent of studies on PFC demonstrating relational structures of task knowledge, such as inferred latent state [42,43], schema [44] and indirectly observed associations between stimuli [45,46].

When moving onto the second delay period during which participants needed to implement the maintained goal on specific memory contents, frontal cortex no longer retained the low-dimensional goal geometry, at least not in an active format, perhaps due to that the frontal 2-D goal geometry was primarily involved in goal maintenance rather than goal implementation. Conversely, the 2-D goal geometry gradually developed in central cortex and later in posterior cortex, indicating that the low-dimensional geometric information was communicated to sensory cortex for goal implementation. Because population-level neural geometry can remain stable when underlying single-neuron activities change [30], one benefit of forming low-dimensional geometry of task information could be that the stable geometry facilitates information relay between cortical regions through communication subspaces [20,21]. This shift in locations was accompanied by a predictability of behavioral performance by posterior goal representations, reminiscent of previous findings showing a relationship between decoding accuracies in visual areas and reaction times using a different task design [14]. Overall, our results are comparable with a growing body of evidence that supports a distributed network for abstract task information during active implementation [16,17], and further extend the idea by proposing that the task information originates from communication between frontal and posterior regions in a low-dimensional format.

In parallel, a similar 2-D stimulus geometry in which color (red-greenness) and size were represented as dissociable dimensions was only found in posterior activities, particularly in primary visual cortex during sample presentation and the ensuing delay. This result lends more credence to our PCA-based approach and aligns with the sensory recruitment account of working memory, which proposes that the storage of sensory stimuli relies on the same visual cortex that initially encode the information [2,3,10]. Besides stimulus-related processing, dedicated sensory regions is also likely to be the locus of goal and stimulus integration, as implementation of task goal and its functional relevance also primarily involved posterior activities.

While the low-dimensional goal representations were present and contributed to memory performance, we acknowledge that other formats can co-exist. Whereas low-dimensional task representations support stability in neuronal readouts, knowledge generalization and efficient learning [18,19], high-dimensional task representations endow high separability between representations and facilitate cognitive flexibility [47]. In fact, these two coding schemes are not mutually exclusive and may be simultaneously present in different brain regions to serve different functional purposes [48]. In our task, goals were implemented once the stimulus appeared, transforming from an abstract concept into an instantiation of concrete state. Therefore, it is possible that high-dimensional codes are also implemented for action selection and output production [19,47,49,50].

### Interaction between goal and stimulus representations in the form of low-dimensional geometry

What mechanisms mediate the transfer of task goals and support the necessary integration of goal and stimulus information? We proposed FMT-to-posterior coherence as a candidate through which higher-order task information is transferred to the locus of specific content storage. More importantly, the transmission retains the geometrical property of the representations, manifested by the relationship between the strength of the connectivity and goal circularity. Although FMT has long been linked to top-down processes during working memory [31,51,52], its functional interpretations are not straightforward [53], ranging from coordinating reactivation of working memory items [31,54], gating of working memory encoding and maintenance [55], to prioritizing internal representation [56]. The evidence provided here points to a more specific role in mediating the communication between frontal and posterior areas for goal representations and the subsequent integration with stimulus representations, while embedded within the general notion that frontal theta oscillation exerts cognitive control signals by synchronizing activity of task-relevant information [57,58]. Furthermore, functional coupling-dependent transmission of goal representations was corroborated by the fMRI connectivity results, involving regions specifically maintained task goals with the matching geometric format. Whilst functional connectomes measured by EEG phase-based metrics and fMRI overlap only to a moderate degree (approximately 0.4) [59], caution is warranted when attributing the cross-modal results to the same underlying neural generator. We nonetheless provide converging support for a fronto-posterior connectivity related to the transmission of two-dimensional goal representations.

Our EEG study also uncovered an interesting dynamic between the representations of remembered and transformed stimuli. Transformed stimulus representations (i.e., participant’s responses) were formed prominently in frontal activities and were accompanied by a decrease in the strength of original stimulus representations. The simultaneous rise of response representation and fall of sample representation, both modulated by fronto-to-posterior coherence, allude to the possibility that stimulus transformation depends on the integration of task goals, and participants might have reduced or even discarded the original copy of remembered items to mitigate interference [60]. However, in the fMRI study, the spatial distinction between the original and transformed stimuli became less evident: representation of transformed stimulus was observed in PFC, whereas those of both original and transformed stimuli were observed in visual cortex. The discrepancy could be caused by the difference in spatial specificities of the two measuring modalities, sensitivity of these signals to stimulus changes, or changes in internal processes induced by the longer delays in the fMRI task. Nevertheless, the observation of response but not stimulus representations in frontal cortex would be consistent with previous studies demonstrating an orthogonal common “template” subspace in PFC ready for response, in contrast to choice-invariant stimulus representations in visual cortex [27]. The two-dimensional response representation in our data might have reflected a similar neural subspace dedicated to guiding behavior. Regarding the variable stimulus representations in visual cortex across experiments, since our study specifically tested for the low-dimensional format which was shown to be functionally relevant to behavior, it is entirely possible that original and transformed stimuli were encoded with lower precision or in a different format [61,62] within specific modalities or ROIs that were not detectable using current methods. As such, the origin of response representations remains to be further tested. Of particular interest, it remains to be explored whether response representation emerges frontally by reading out the input from goal and stimulus interactions in posterior cortex, or it is formed posteriorly instead and subsequently transmitted to frontal cortex.

### Differential functions of DLPFC and MPFC

The limited spatial resolution of EEG has precluded precise localization of the two-dimensional goal geometry in frontal cortex. With the follow-up fMRI experiment, we further localized the goal geometry to subregions within DLPFC and MPFC. Specifically, the DLPFC cluster centers on the superior frontal gyrus (SFG) and lies superior to the conventional Area 9-46. Consistent with the hypothesized origin of midline theta activity in EEG [53,63], both regions lie close to the frontal midline, with one to the more lateral side and the other to the more medial side. This anatomical overlap points to an intriguing possibility that the two identified PFC subregions with significant goal geometry likely relate to the source of the cognitive control signals driving FMT. Surprisingly, although both subregions demonstrated a two-dimensional goal geometry, only the functional connectivity between SFG and visual cortex predicted the goal geometry in the latter during later delay. This functional distinction between DLPFC and MPFC aligns with recent theoretical work on the differential roles of task representations in lateral and medial PFC, purporting that the LPFC uses task representations to build rules for action selection, whereas the MPFC abstracted task knowledge in a relational map [41]. In line with this notion, only DLPFC is actively involved in the coordination between frontal and visual cortex which likely supports the transmission of goal geometry for implementation. Our newly designed paradigm offers a useful tool for investigating the functional distinction between the two networks. Future work with a more comprehensive examination on the neural geometries in subregions of the two networks is needed to further address this question.

## Conclusion

In summary, across two experiments, we provided converging evidence for how distributed task representations emerge and transfer in working memory to support goal-directed behaviors. In particular, frontal cortex maintains task-specific, low-dimensional neural geometries of goal representations in preparation for subsequent goal implementation in posterior visual cortex. Our findings highlight working memory as a multi-component and collaborative cognitive system that relies on coordinated neural interactions across multiple brain regions.

## Materials and Methods

### Participants

Twenty-three participants were recruited for the EEG experiment (mean age = 24.3 years; age range = 19-30 years; 14 females), one was excluded due to excessive noise (over 20% of total trials). Twenty-one MRI-eligible participants were recruited for the fMRI experiment (mean age = 24.2 years; age range = 21-30 years; 16 females). There is no overlap in participants between studies. All participants were recruited from the Shanghai Institutes for Biological Sciences community, reported neurologically and psychologically healthy, had normal or corrected-to-normal vision, provided written informed consent, and were monetarily compensated for their participation. The study was approved by the Ethics Committee of the Center for Excellence in Brain Science and Intelligence Technology, Chinese Academy of Sciences (CEBSIT-2020028).

### Experimental design and procedure

#### Overview

We designed an experimental task that required the maintenance of both a goal and a specific stimulus in working memory and at later stage the manipulation of the stimulus features based on the goal. Specifically, each remembered stimulus varied along two orthogonal stimulus dimensions, size (small to big) and color (green to red). Correspondingly, there were totally four types of goals composited by two orthogonal dimensions of size and color manipulations: adjusting the remembered stimulus to be **1)** bigger and redder, **2)** bigger and greener, **3)** smaller and redder, and **4)** smaller and greener. The two-dimensional stimulus space, as well as the extent of required adjustment in size and color were predefined and learned in a behavioral session preceding the main task.

#### Definition and manipulation of stimulus features

Color (green-redness) were adjusted by first converting the images into grayscale to remove all original hues while preserving the opacities of pixels using the Image module from Python package PIL. Then the resulting files were converted to RGBA mode again and the green-redness values were manipulated by adding/subtracting the same value in the Red and Green channels whereas the Blue channel was set to zero. This made so that the stimulus transitioned from extremely green to extremely red in a smooth fashion (for illustration see Figure 1D). The acceptable RGB value range for PsychoPy was 0-1, each button press moved the R and G value of all image pixels to opposite directions by 0.01. The values in R and G channels were rescaled to 0.25-0.75 with a mean of 0.5. Therefore, for each stimulus there existed 150 variations of color-manipulated images (until all pixels had a value of 0 or 1). However, since all pixels appearing uniformly green or red would render the stimulus unidentifiable (because there is no contrast), the 30 images with most extreme R/G values were discarded, resulting in 120 steps for the range of color adjustment. Both starting size and colors were drawn from three predetermined bins, resulting in nine unique conditions, which were used for subsequent analyses. For size the bins were 0.17±0.01, 0.22±0.01 or 0.27±0.01 of screen height; for color they were 34±2, 58±2 and 82±2, indexing from the 120 color steps. Stimulus size was directly controlled in PsychoPy program using the size attribute and the allowed range was set to 0.01-0.45 screen height. The distance between each particular sample stimulus and its target answer for size was ±24% of the original size (unit: screen height), and ±26 (unit: color steps) for color. Participants learned the required distance during a behavioral learning session.

#### Behavioral learning

In the behavioral session, which was held one or two days before the main task session, participants learned the degree of required adjustment by first viewing all pairs of starting and target values in size and color shown side-by-side, respectively (162 trials per stimulus feature), before receiving a Two-Alternative-Force-Choice (2-AFC) test whereby they were given a starting stimulus and asked to choose the correct target stimulus from two options (80 test trials per stimulus feature). Next, both features were combined together in the same stimulus in another round of 2-AFC test to familiarize them for simultaneous adjustment of both size and color based one of the four goals, in preparation for the main working memory task (100 trials). Finally, participants completed 180 trials for the main task in order to apply the learned degree of adjustment on stimulus features. In the EEG experiment, the four goals were associated respectively with different shape cues, which the participants also learned and practiced the association between shape cues (circle, square, triangle and pentagon; Figure 1A) and goals in these 180 trials. In the behavioral session only, trial-wise feedback of response error was given so that participants could keep improving their performance.

#### EEG main task

The main task consisted of two memory delays and a response period. The first delay required the maintenance of the goal only, and the second delay required both the maintenance of the goal and the stimulus. At the beginning of a trial, there was a 300-ms fixation period followed by one of the shape cues for 400 ms appearing centrally (Goal cue), signaling the direction of manipulation in the current trial. This was followed by a 1300-ms delay (Delay 1), during which participants should keep maintaining the manipulation goal. Following the first delay, a stimulus was presented for 600 ms (Sample). The stimuli were randomly chosen from the exemplars belonging to three different conceptual categories with similar shapes: bowling pins, plants and microphones, provided and detailed in [64]. Participants were instructed to maintain the sample’s size and color during a second delay period for 1500 ms (Delay 2). To prevent participants from using physical changes on the retinal image as memory aids, the stimulus was presented in a random location of an invisible circle around the center of the screen with a radius of 0.08 screen height, making it a 2° visual angle difference from the center. During the response period (Response), the same stimulus reappeared with a randomly chosen but different size and color from the remembered values to prevent response preparation. Participants were given a maximum of 6000 ms to adjust the features of stimulus on screen to the correct degree using button presses. The correct response depended on both the starting values and the cued goal, as learned in the behavioral session. Thereafter, a variable intertrial interval was incurred (800-1200 ms, drawn from a uniformed distribution). Overall, participants completed 648 trials divided into 12 blocks lasting for ~2 h, with task variables of goal types and stimulus features counterbalanced, resulting in 18 trials in each of the possible combinations. Of note, during the EEG recording participants only received averaged feedback on their performance after a block, to avoid further improvement on the extent of change per se as a result of trial-wise feedback.

#### fMRI main task

Behavioral paradigm and experimental procedure in the fMRI experiment generally remained consistent with the EEG study unless otherwise stated. In-scanner task shared the same components as in EEG but with adjusted length to accommodate the delay in hemodynamic response function (Figure 1E bottom). The durations of epochs were as follows: Goal cue = 500 ms; Delay 1 = 5500 ms; Sample = 600 ms; Delay 2 = 8400 ms; Response = 5000 ms. Intertrial intervals were randomly chosen from 4000, 5500 and 7000 ms with equal likelihood. Of note, goal cue was presented as texts instead of associated shapes in the fMRI task, therefore participants were no longer required to learn the pairings. In total participants completed 12 functional blocks, each containing 18 trials and lasting 466.5 s.

#### EEG Apparatus

Stimulus presentation was implemented using PsychoPy (version 2021.2.3) [65] on a 48 × 27 cm HIKVISION LCD screen with a 60 Hz refresh rate and a 1920 × 1080 resolution. Stimuli were shown in white font on a gray background (RGB = 128, 128, 128) at a distance of 62 cm. During the task, head position was stabilized by a chin rest. Responses were given with the right hand on the four arrow keys on a keyboard. EEG data were recorded using a Brain Products ActiCHamp recording system and BrainVision Recorder Software (Brain Products GmbH, Gilching, Germany). Scalp voltage was obtained from a broad set of 59 channels at 1000 Hz according to the extended 10-20 positioning system (FCz as reference). Channel impedance was kept below 20 kΩ.

#### EEG preprocessing

EEG data were preprocessed in MNE-Python [66], which firstly involved down-sampling to 250 Hz and bandpass filtering using a high-pass filter of 0.1 Hz and a low-pass filter of 40 Hz. The continuous raw data were then segmented into epochs, corresponding to 500 ms before the onset of the goal cue (200 ms before the fixation onset) until 600 ms after the onset of the response period. EEG channels with excessive noise were identified through visual inspection and replaced via interpolation using a weighted average of the surrounding channels. Each epoch was inspected visually for artifacts such as excessive muscle movements and amplifier saturation, and contaminated trials were discarded. Stereotyped artifacts such as ocular movements were subsequently removed from the data via independent component analysis. The data were baseline corrected for the subsequent analyses using signals from the time window of −200 to 0 ms before fixation onset. There were on average 621 trials per person remained after epoch rejection (SD = 26 trials).

#### fMRI Data Acquisition

MRI scanning was performed at the Functional Brain Imaging Platform (FBIP), Center for Excellence in Brain Science and Intelligence Technology (CEBSIT), on a Siemens 3T Tim Trio MRI scanner with a 32-channel head coil. High-resolution T1-weighted anatomical images were acquired using a magnetization-prepared rapid gradient-echo (MPRAGE) sequence (2300 ms time of repetition (TR), 3 ms time of echo (TE), 9° flip angle (FA), 256 × 256 matrix, 192 sequential sagittal slices, 1 mm^3^ isotropic voxel size). Whole-brain functional images were acquired using a multiband 2D gradient-echo echo-planar (MB2D GE-EPI) sequence with a multiband acceleration factor of 2, 1500 ms TR, 30 ms TE, 60° FA within a 74 × 74 matrix (46 axial slices, 3 mm^3^ isotropic voxel size). After four experimental runs we also acquired whole-brain Fieldmap phase and magnitude images for correction of EPI distortions. Stimuli were presented on a 1280 × 1024 resolution MRI-compatible screen at the back of the scanner, and participants viewed the screen through a mirror attached to the head coil with a viewing distance of 90.5 cm. They used two two-button response boxes, one in each hand to adjust the stimuli.

#### fMRI Preprocessing

Preprocessing of MRI data was performed using fMRIPrep 21.0.2 [67], which is based on Nipype 1.6.1 [68]. The single-band reference data were taken as the reference volume of each run, and were co-registered to the anatomical scan. Both the anatomical and functional scans were normalized to the MNI152 template. A detailed description of preprocessing procedure can be found in Supporting information.

### Quantification and statistical analyses

#### EEG: Neural geometries of task representations

To uncover the geometry of neural representations associated with goals, sample stimuli and responses, and describe their similarities to goal/stimulus space, we conducted principal component analysis (PCA) to identify low-dimensional spaces in which the relevant task variables were encoded, respectively. Specifically, each participant’s voltage data was collapsed over trials from the same conditions (goals, stimulus or response bins), with each column standardized independently before applying PCA, resulting in a matrix of shape n_conditions × n_channels. Note that we used the same standard to divide sample stimuli and responses into bins, as the latter are essentially manipulated stimuli that share the same feature value ranges as the former. We repeated this procedure for frontal and posterior channels separately, within each temporal epoch of a trial (i.e., Goal cue: 0-400 ms; Delay 1: 400-1700 ms; Sample: 1700-2300 ms; Delay 2: 2300-3800 ms; and Response (3800-4300 ms) during which the data was averaged. Frontal channels include: Fp1, Fz, F3, F7, F4, F8, Fp2, AF7, AF3, AFz, F1, F5, F6, AF8, AF4, F2, and posterior channels include: Pz, P3, P7, O1, Oz, O2, P4, P8, P1, P5, PO7, PO3, POz, PO4, PO8, P6, P2. Unless stated otherwise, the PCA-related analyses and ensuing behavioral correlations were based on PCA trained on these individual data matrices. For the group-based visualization of the subspace, the individual averaged activity patterns were subsequently concatenated horizontally, resulting in a matrix of shape n_conditions × (n_channels × n_participants). All remaining steps followed the aforementioned procedure for individual data.

#### EEG: Circularity index

To quantitively assess the structure of representations in the space reconstructed by PCA in spite of differences between task variables, individuals or principal planes, we used a simple metric, namely circularity index to capture the spatial properties of the neural geometry. Briefly put, circularity [69] of a shape was defined as the ratio between its area and the square of the total length of perimeter:

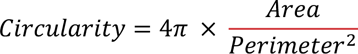

A circle would always have a circularity of 1. If goal representations abided the relational structure formed in the hypothetical goal space consisting of two orthogonal dimensions (goal size and goal color) with equal distance between goals, the circularity would be that of a square which is ~0.78. Since there were only four points to define the structure, this value was also the maximum as any quadrilateral would have a smaller circularity. To calculate circularity in the neural space, borders were drawn between the projected coordinates of each unique conditions, in the same order as in the hypothetical space (Figure 1B). For instance, for goal representations, the points were connected in order of 1 (bigger & redder) – 2 (bigger & greener) – 4 (smaller & greener) - 3 (smaller & redder). Area and perimeter were then calculated using Python package Shaply.

The calculation of circularity index for sample stimuli and responses was similar to that of the goal space except for there were nine conditions/points to consider (three bins per stimulus feature). For the sake of constructing a close-formed geometric shape within the neural subspace for which circularity index could be calculated, the medium color value was not used, leading to six effective coordinates in the stimulus feature space.

To test the significance of circularity index values, trial labels were shuffled within participants before repeating the above PCA and circularity analyses for 5000 times. The true values were compared to the resulting null distribution to derive *p*-values.

#### EEG: Cross-validated time course of circularity index

To investigate the temporal relations of 2-D goal geometry across groups of channels, we calculated the cross-validated circularity time courses. Instead of averaging data within each trial epoch, data was averaged within an 80-ms sliding window which were non-overlapping, resulting in 60 points per trial). Each individual’s trial-wise data was bootstrapped for 2000 times in a stratified manner, which subsequently went through the same group-level PCA procedure described above. Additionally, for each iteration of the bootstrapping, a baseline was also achieved through shuffling the resulting trial labels 20 times and taking the means of circularity indices. The difference between the resulting circularity time series from bootstrapped and shuffled data were taken to perform a cluster-based permutation test (MNE-Python *permutation_cluster_1samp_test*) to determine significance. In this section of analysis, the central channels were included, which consisted of the following electrodes: FC5, FC1, C3, T7, CP5, CP1, CP6, CP2, Cz, C4, T8, FC6, FC2, FT7, FC3, C1, C5, TP7, CP3, CPz, CP4, TP8, C6, C2, FC4, FT8.

#### EEG: behavioral relevance analysis based on trial splitting

For goal geometry, the bottom and top quartiles (i.e., 25 percentage) of each participant’s trial-wise data were taken based on the magnitude of combined response error to compute group-level circularity indices corresponding to the good and bad trials within each epoch, respectively. The two sets of data were further bootstrapped for 2000 times and within each iteration, the group circularity index for each quantile was calculated separately (i.e., individuals’ data were concatenated horizontally before being projected to the subspace identified by a PCA trained on the whole dataset, thus trials from both quartiles were compared on an equal ground) and the difference was taken. The resulting distribution of differences was compared against zero and p-values were derived by counting the proportion of values below zero according to a one-tailed hypothesis of 2-D goal geometry of bottom quartile (i.e., smaller errors) being stronger than that of the top one. For stimulus-specific geometry, we instead median-split the trials due to the quartile approach could not cover all unique conditions needed for the calculation.

#### EEG control analysis: Time-resolved representational similarity analysis (RSA)

To validate the goal representations in the 2-D space revealed by PCA, we additionally performed an RSA analysis [70] on the individual level. For each participant, we tested whether a two-dimensional RSA model of task goals would explain the data. This 2-D RSA model assumed that the pairwise representational distances between goals should resemble the 2-D geometry as revealed by PCA (Figure 3). Specifically, the model representational dissimilarity matrix (RDM) for 2-D goals was constructed by counting the number of same values across the two goal dimensions (i.e., the hamming distance: 0 for same goals, 1 for matching on one dimension but different on the other, and 2 for two goals with non-overlapping values). We included both task goals and stimulus features to define unique conditions in order to enrich the number of condition pairs, as with only the binary goal dimensions, number of valid pairs would be too low to allow stable estimates of model-neural correlations. For data RDM, cross-validated (4 folds) cosine distance was computed from the standardized data, averaged within an 80-ms non-overlapping sliding window. Comparison between the neural and model RDMs was performed using MNE-Python’s function *mne_rsa.rsa* and Spearman rank correlation as metric. Significance of RSA time courses from all individuals were tested against 0 using MNE-Python function *permutation_cluster_1samp_test*.

#### EEG control analysis: Motor-related neural geometry

To ensure that the 2-D goal geometry was not caused by signals related to motor preparation, a series of control analyses were conducted. The motor-specific conditions were defined by subtracting the initial values of the response object (which were randomly drawn for a uniform distribution) from the values of the final response. The signs of the resulting feature differences indicated the directions of adjustment the participants performed on that trial (e.g., to make the object bigger and redder from the initial status, one needed to press the buttons corresponding to “bigger” and “redder”), which were taken as the trial label specific to motor preparations. The following steps to calculate group-based circularity index (for correlation with goal-specific circularity) and cross-validated circularity time course (for significance testing) were all identical to the procedures stated above. Chi-square tests for independence between possible motor preparation signals and task goals, and between stimulus and task goals, were performed using the Scipy library.

#### EEG: Coherence analysis

To understand how frontal control signals modulated goal-related representations, we calculated a coherence measure to investigate time-resolved connectivity between frontal and posterior channels. In particular, frontal medial theta oscillation (FMT) has been considered to be the medium through which executive control mechanisms orchestrate working memory contents [53,63]. Moreover, goal representations could serve as the signals that modulate specific items stored in working memory, as observed in various studies that abstract task information were present in prefrontal regions [14,15,37]. To this end, coherence values were computed across all time points with the MNE-Python *spectral_connectivity_epochs* function, which involved calculating the cross-spectral densities, and estimating pairwise coherence in the range of 4-7 Hz between the seed frontal channel (Fz) and every posterior channel before being averaged. We chose the metric of weighted phase lag index (wPLI) as it is more robust against artifacts dues to volume conduction and noise [71]. Significant clusters in the baseline-corrected coherence values were determined using the MNE-Python function *permutation_cluster_1samp_test* (one-tailed). To test whether significant FMT clusters during delay were functionally related to strength of representational geometries, the significant group clusters were taken as frequency and temporal masks to select coherence values from every individual’s result, which were correlated with the circularity indices at each epoch of the trials, respectively.

#### FMRI: General linear models (GLM)

We fit a single-trial level univariate GLM to extract neural response estimates for each delay period. For each functional run, the design matrix included task regressors representing the separate periods of a trial and trial-wise regressors associated with the period of interest. For example, for modelling beta series of the first delay, the task regressors would consist of Goal cue (500 ms), Sample (600 ms), Delay 2 (8400 ms), Response (5000 ms), and trial-wise regressors for Delay 1 (i.e., Delay1_trial1(5500 ms), Delay1_trial2, etc). The procedure was repeated for other time periods of interest. Thus, each trial was separated out into its own condition within the design matrix [35]. Additionally, the model also included six head-motion regressors, three global signals from CSF, white matter and whole-brain, and three trend predictors from a polynomial drift model. Task regressors were convolved with the SPM canonical hemodynamic response function (HRF) and its time derivative. Functional data were standardized, high-pass filtered at 0.01 Hz and spatially-smoothed with a 6-mm FWHM kernel. The resulting trial-wise beta series were brought forward to subsequent analyses.

For selection of 500 most activated voxels in each ROI, a generic GLM was fitted to individual BOLD signals in which each task period was modeled as a unique regressor. All other settings were the same as in the single-trial GLM described above. For each participant, the estimated coefficients were used to select within ROIs the top 500 most positively activated voxels to repeat the PCA-based neural subspace analysis. For goal geometry calculation, we used voxels selected based on the Delay 1 and Delay 2 regressors, respectively. For sample geometry calculation, we used voxels selected based on the Sample and Delay 2 regressors, respectively.

#### FMRI: Region of interest (ROI) analyses

To localize brain regions with the same neural geometries as identified in the EEG results more specifically, we defined subregions in the frontal and posterior visual cortex before replicating the analysis steps for identifying representational structures. These anatomical ROIs were selected based on previous work using a multi-modal cortical atlas [32]. For frontal regions, we selected section 19 (anterior cingulate and medial prefrontal cortex), 20 (orbital and polar frontal cortex), 21 (inferior frontal cortex) and 22 (dorsolateral prefrontal cortex) as referred in the original paper, which were each further divided into two separate parcels as the entire area was relatively large (Figure 5). For posterior regions, the following visual-related sections were chosen: 1 (primary visual cortex, V1), 2 (early visual cortex/extrastriate cortex), 3 (dorsal visual cortex), 4 (ventral visual cortex) and 5 (MT complex and neighboring visual areas). All ROIs were merged bilaterally for the ROI-based PCA analysis, and were left unilaterally for the connectivity analysis.

The bilateral ROI masks were used to select voxels from the trial-wise coefficient maps before applying the same procedures of calculating PCA and circularity index. Furthermore, to test the stability of results derived from these ROIs, the analyses were repeated on the 500 most positively activated voxels within each ROI, separately defined for each participant using task effects corresponding to the trial epochs (e.g., Delay 1 & Delay 2), estimated using GLMs.

#### FMRI: ROI-based neural geometries of task representations

Consistent with the EEG analyses, for group-level visualization and testing for significant 2-D geometry, averaged beta coefficients for each unique conditions (goals or stimulus bins) within an ROI were concatenated across participants to form a matrix of shape n_conditions × (n_voxels × n_participants) before PCA was applied. Individual matrices of shape n_conditions x n_voxels were used instead for correlational analyses.

#### FMRI: Beta series functional connectivity

Following the result of ROI-based geometries, inter-regional connectivity between ROIs was investigated. The beta values corresponding to both delay periods estimated from the trial-wise GLM procedure described above were entered into a partial correlation estimation using Nilearn’s *ConnectivityMeasure* class. Specifically, for all unilateral ROIs, parameter estimates were averaged within each ROI and partial correlation coefficients were calculated between every pair of ROIs, taking into account the influence of all other regions. For statistical testing, we applied Fisher’s z transformation on the correlation coefficients and conducted one-sample t-test against 0 (α=0.05, Bonferroni corrected). We also examined whether the unilateral functional connectivity between a frontal and a posterior ROI was predictive of circularity indices of 2-D representational geometries in the latter, as well as behaviors using Spearman rank correlation (one-tailed).

#### FMRI control analysis: RSA

Similar to the EEG RSA analysis, an RSA was conducted on the trial-wise delay-period beta estimates to offer complementary evidence for the PCA analyses. Definition of the model RDM was the same as used in EEG; for neural data RDM, cross-validated (4 stratified folds) cosine distance was computed from the standardized data. Comparison between the neural and model RDMs was performed using MNE-Python’s function *mne_rsa.rsa* and Spearman rank correlation as metric. Significance of correlations were tested with a one-sided t-test against 0.

## Supplemental Figures

**Figure S1.**
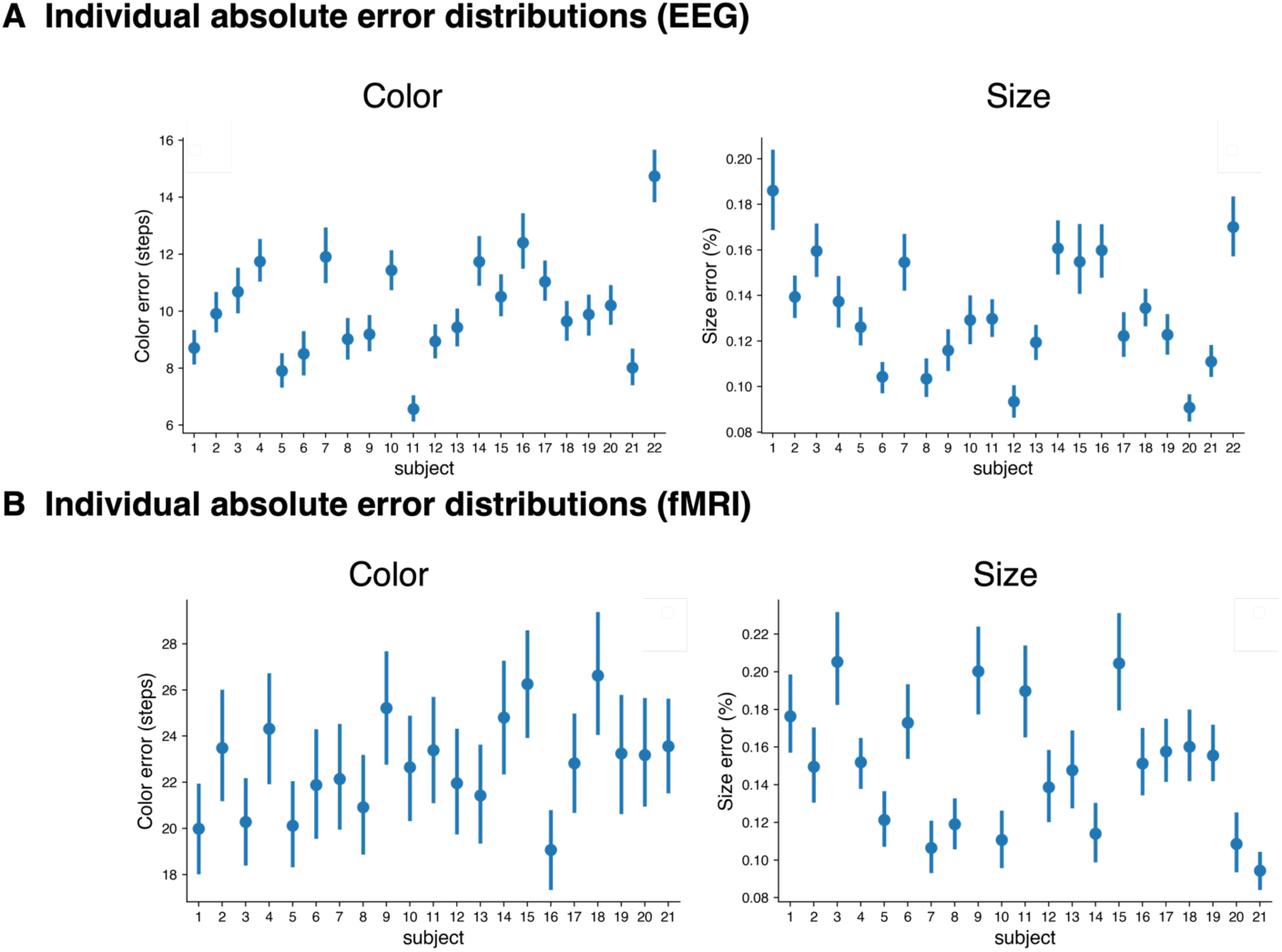
Absolute error distributions for each participant. Error bar represents 95% confidence interval.

**Figure S2.**
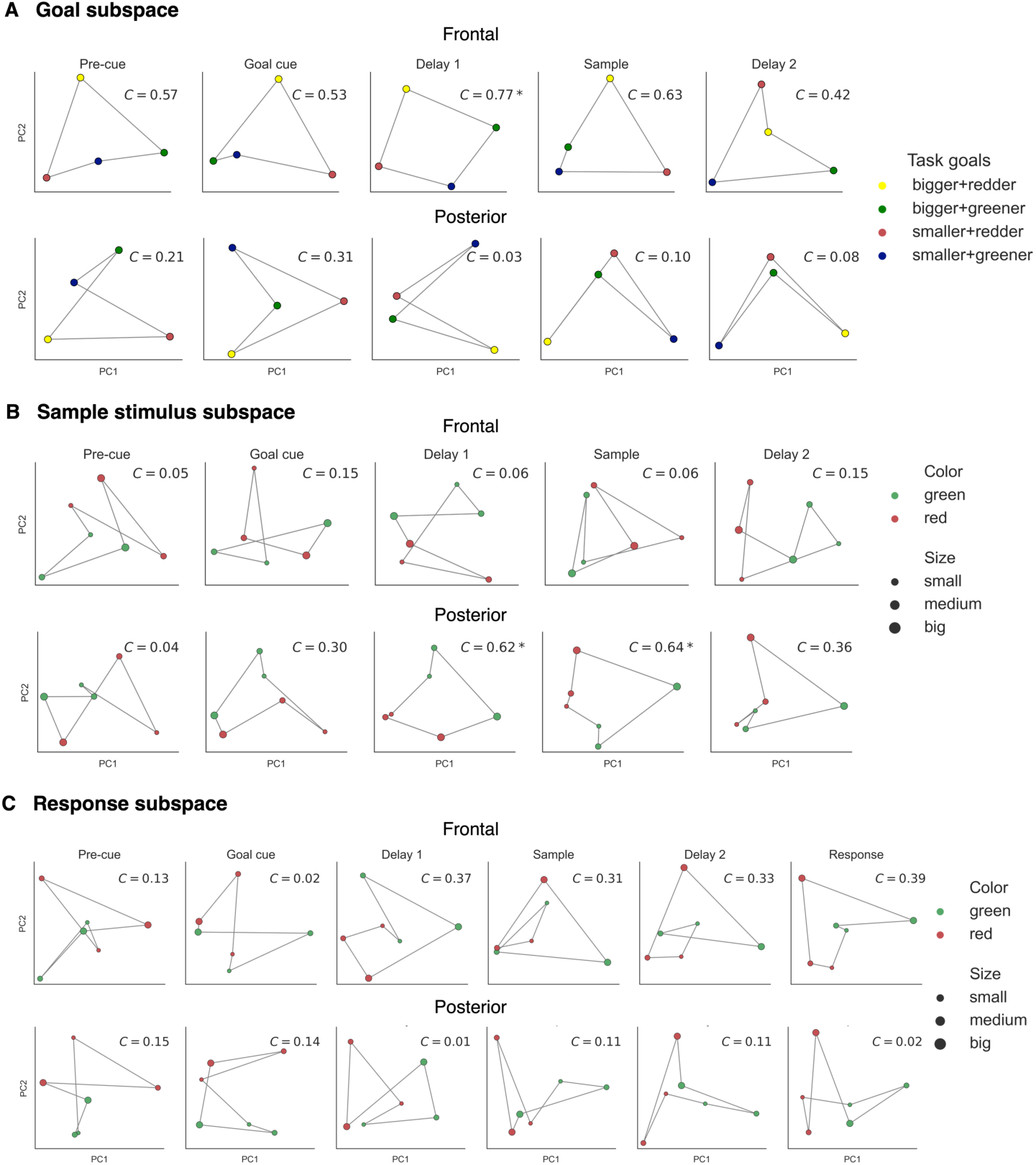
Neural geometries using averaged data within each task epoch **(A)** Representational structures in the identified task goal subspace for frontal (upper) and posterior (lower) activities. Each colored dot represents a unique condition. *C* denotes circularity index and asterisk denotes significance from permutations (α = .05). **(B)** Same as **(A)** but for stimulus geometry (using six conditions). **(C)** Same as **(B)** but for response geometry, using participants’ answers as feature values.

**Figure S3.**
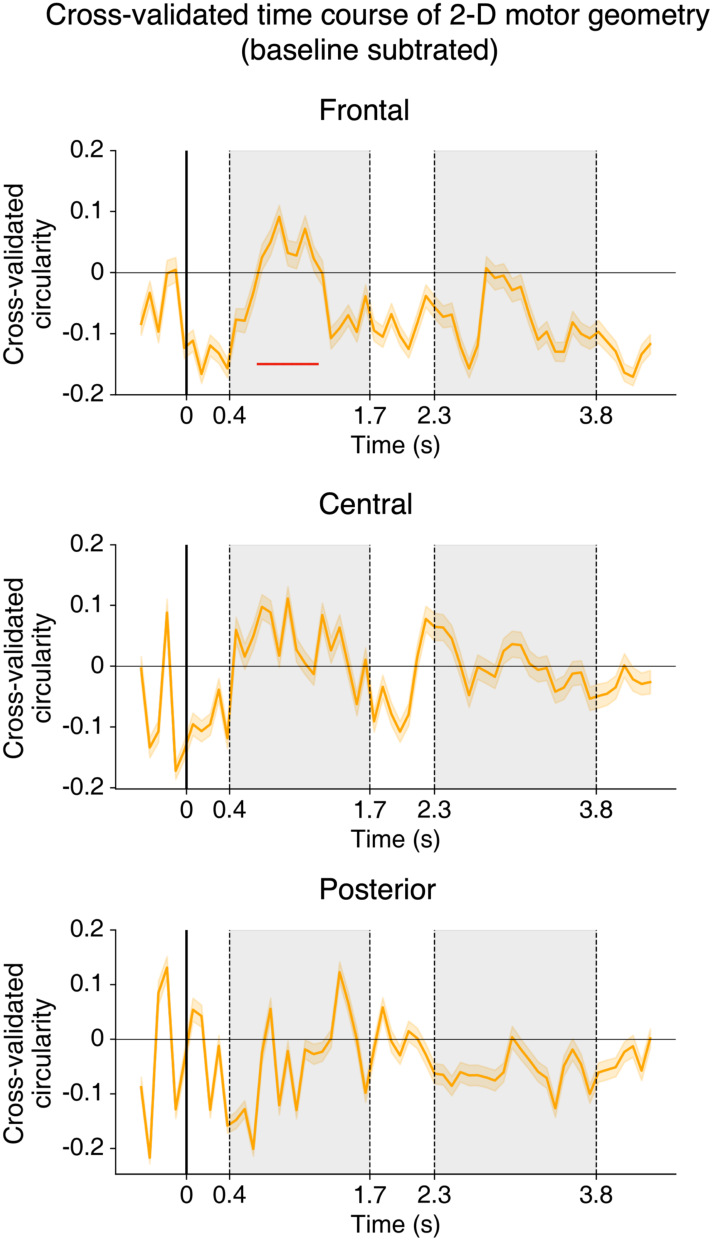
Circularity time courses of motor preparation signals for frontal, central and posterior channels. Red horizontal lines denote significant time points (α = .05) corrected using a cluster-based permutation test.

**Figure S4.**
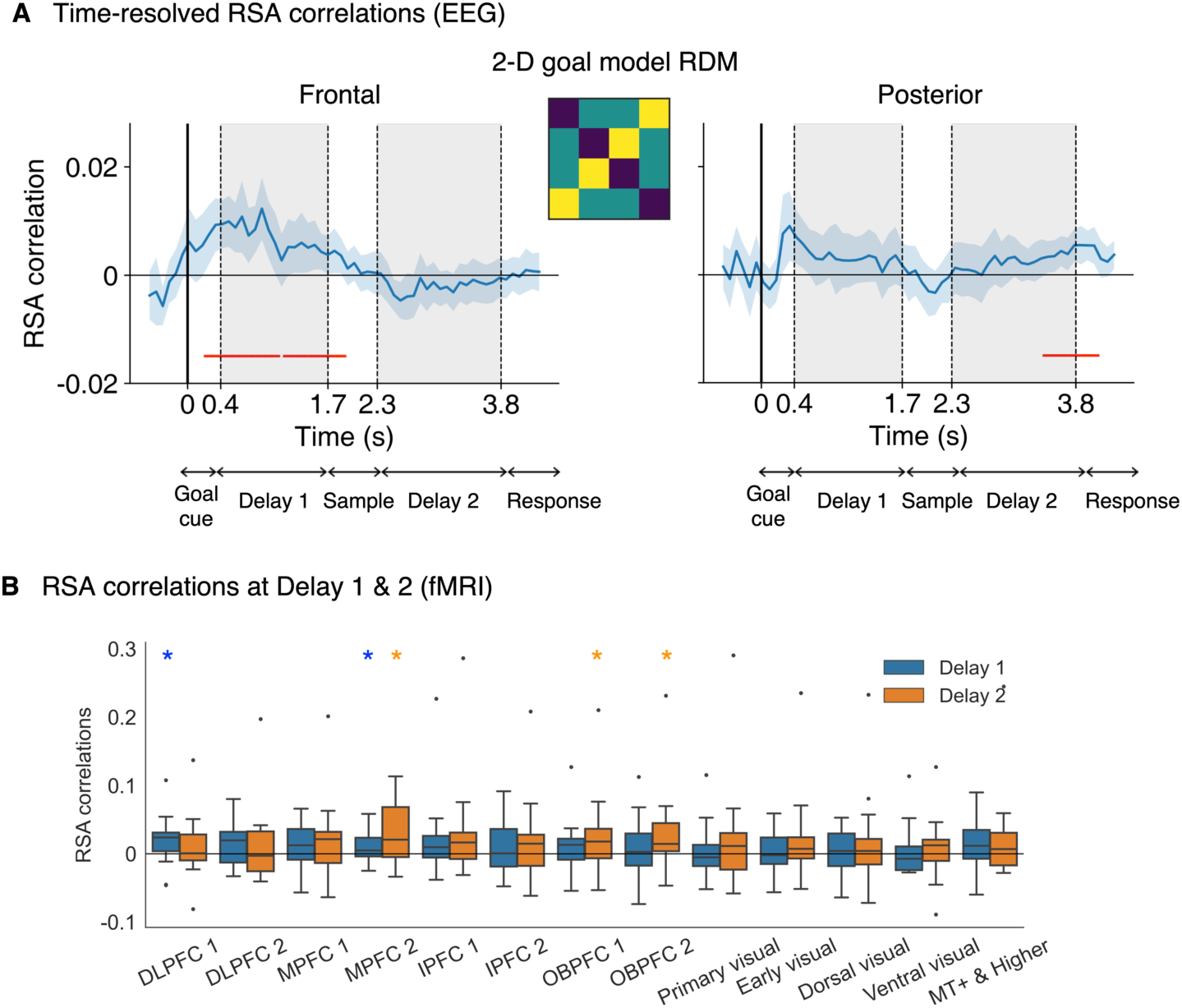
Representational similarity results for EEG and fMRI data. **(A)** Time-resolved cross-validated RSA correlations with individual condition-averaged data using the 2-D goal model in frontal and posterior channels, respectively. Data was averaged temporally within a non-overlapping 80-ms sliding window. Red horizontal lines denote significant time points (α = .05) corrected using a cluster-based permutation test. Error bars represent 95% confidence interval. **(B)** Cross-validated RSA correlations using beta coefficients for Delay 1 and 2. Asterisks denotes *p* < 0.05 (uncorrected) in a one-sided t-test against 0.

**Figure S5.**
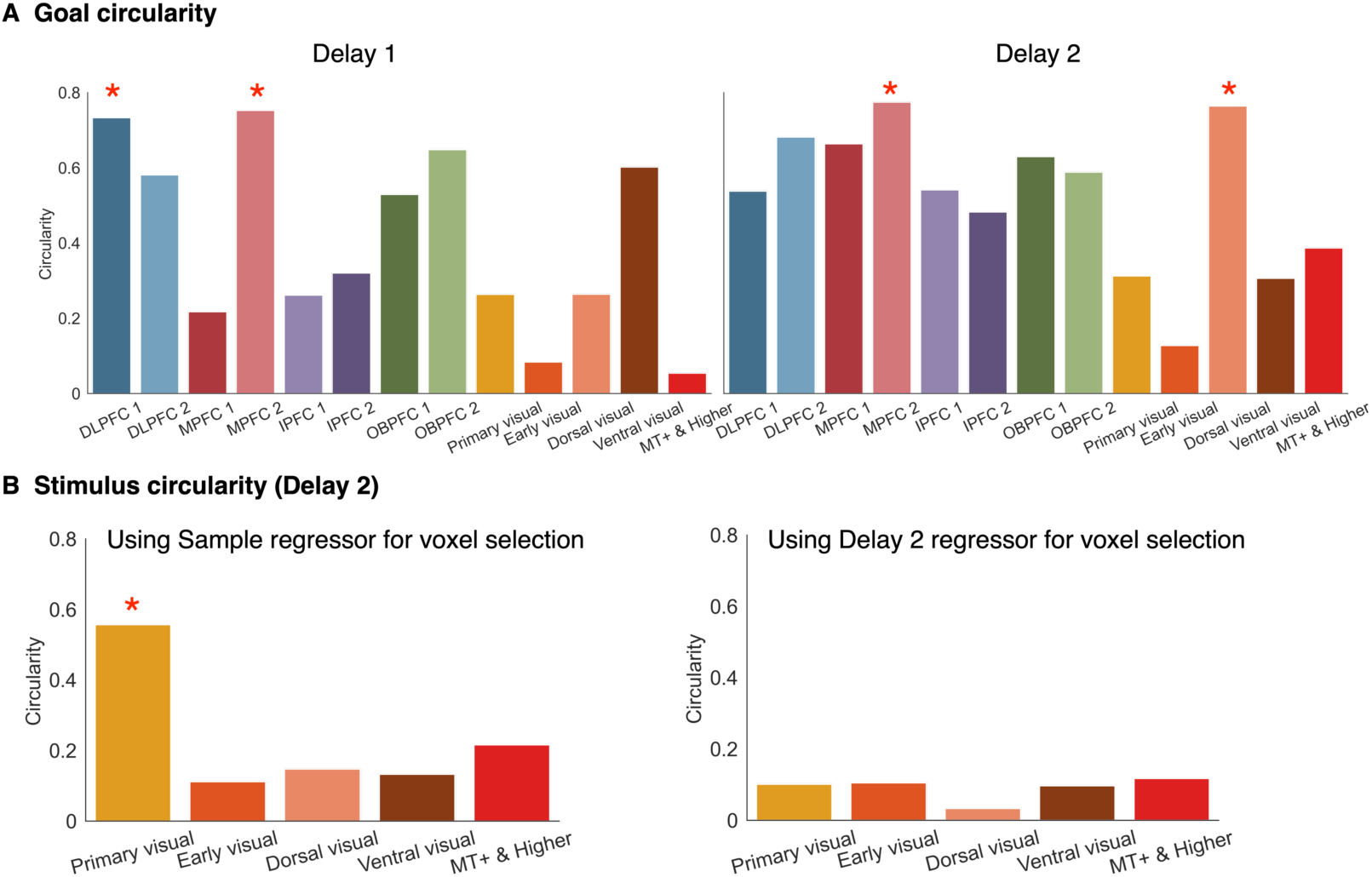
FMRI ROI-based neural geometry results using 500 most positively activated voxels. **(A)** Goal circularities for all PFC and visual ROIs. Voxels were selected using task regressors for Delay 1 and 2 respectively, for each individual. Red asterisk denotes *p* < 0.05 based on permutations. **(B)** Same as **(A)** but for stimulus circularity. The result using voxels selected with the Sample regressor (i.e., BOLD activities were highest during Sample presentation) were consistent with our primary result using entire ROIs, but not when using the Delay 2 regressor.

**Figure S6.**
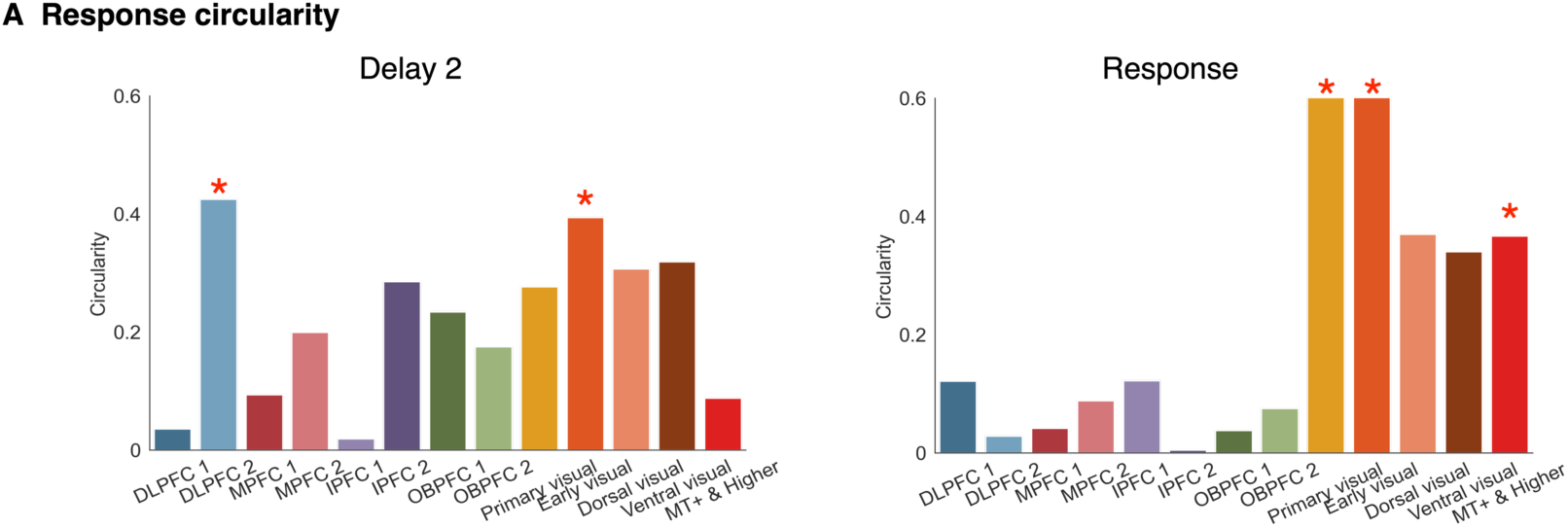
(A) FMRI ROI-based neural geometry for response representation. Red asterisk denotes *p* < 0.05 based on permutations.

## References

1. Baddeley A. Working memory: looking back and looking forward. Nat Rev Neurosci. 2003;4(10):829–39. doi: 10.1038/nrn1201.

2. Harrison SA, Tong F. Decoding reveals the contents of visual working memory in early visual areas. Nature. 2009;458(7238):632–5. doi: 10.1038/nature07832.

3. Serences JT, Ester EF, Vogel EK, Awh E. Stimulus-specific delay activity in human primary visual cortex. Psychol Sci. 2009;20(2):207–14. doi: 10.1111/j.1467-9280.2009.02276.x.

4. Riggall AC, Postle BR. The Relationship between Working Memory Storage and Elevated Activity as Measured with Functional Magnetic Resonance Imaging. Journal of Neuroscience. 2012;32(38):12990–8. doi: 10.1523/Jneurosci.1892-12.2012.

5. Emrich SM, Riggall AC, Larocque JJ, Postle BR. Distributed patterns of activity in sensory cortex reflect the precision of multiple items maintained in visual short-term memory. J Neurosci. 2013;33(15):6516–23. doi: 10.1523/JNEUROSCI.5732-12.2013.

6. Ester EF, Sprague TC, Serences JT. Parietal and Frontal Cortex Encode Stimulus-Specific Mnemonic Representations during Visual Working Memory. Neuron. 2015;87(4):893–905. doi: 10.1016/j.neuron.2015.07.013.

7. Gosseries O, Yu Q, LaRocque JJ, Starrett MJ, Rose NS, Cowan N, et al. Parietal-Occipital Interactions Underlying Control- and Representation-Related Processes in Working Memory for Nonspatial Visual Features. J Neurosci. 2018;38(18):4357–66. doi: 10.1523/JNEUROSCI.2747-17.2018.

8. Yu Q, Shim WM. Occipital, parietal, and frontal cortices selectively maintain task-relevant features of multi-feature objects in visual working memory. Neuroimage. 2017;157:97–107. doi: 10.1016/j.neuroimage.2017.05.055.

9. Yu Q, Shim WM. Temporal-Order-Based Attentional Priority Modulates Mnemonic Representations in Parietal and Frontal Cortices. Cereb Cortex. 2019;29(7):3182–92. doi: 10.1093/cercor/bhy184.

10. D’Esposito M, Postle BR. The cognitive neuroscience of working memory. Annu Rev Psychol. 2015;66:115–42. doi: 10.1146/annurev-psych-010814-015031.

11. Bunge SA, Kahn I, Wallis JD, Miller EK, Wagner AD. Neural circuits subserving the retrieval and maintenance of abstract rules. J Neurophysiol. 2003;90(5):3419–28. doi: 10.1152/jn.00910.2002.

12. Harel A, Kravitz DJ, Baker CI. Task context impacts visual object processing differentially across the cortex. Proc Natl Acad Sci U S A. 2014;111(10):E962–71. doi: 10.1073/pnas.1312567111.

13. Wallis JD, Anderson KC, Miller EK. Single neurons in prefrontal cortex encode abstract rules. Nature. 2001;411(6840):953–6. doi: 10.1038/35082081.

14. Muhle-Karbe PS, Duncan J, De Baene W, Mitchell DJ, Brass M. Neural Coding for Instruction-Based Task Sets in Human Frontoparietal and Visual Cortex. Cereb Cortex. 2017;27(3):1891–905. doi: 10.1093/cercor/bhw032.

15. Woolgar A, Thompson R, Bor D, Duncan J. Multi-voxel coding of stimuli, rules, and responses in human frontoparietal cortex. Neuroimage. 2011;56(2):744–52. doi: 10.1016/j.neuroimage.2010.04.035.

16. Gonzalez-Garcia C, Formica S, Wisniewski D, Brass M. Frontoparietal action-oriented codes support novel instruction implementation. Neuroimage. 2021;226:117608. doi: 10.1016/j.neuroimage.2020.117608.

17. Formica S, Gonzalez-Garcia C, Senoussi M, Marinazzo D, Brass M. Theta-phase connectivity between medial prefrontal and posterior areas underlies novel instructions implementation. eNeuro. 2022;9(4). doi: 10.1523/ENEURO.0225-22.2022.

18. Mack ML, Preston AR, Love BC. Ventromedial prefrontal cortex compression during concept learning. Nat Commun. 2020;11(1):46. doi: 10.1038/s41467-019-13930-8.

19. Badre D, Bhandari A, Keglovits H, Kikumoto A. The dimensionality of neural representations for control. Curr Opin Behav Sci. 2021;38:20–8. doi: 10.1016/j.cobeha.2020.07.002.

20. Semedo JD, Zandvakili A, Machens CK, Yu BM, Kohn A. Cortical Areas Interact through a Communication Subspace. Neuron. 2019;102(1):249–59 e4. doi: 10.1016/j.neuron.2019.01.026.

21. Srinath R, Ruff DA, Cohen MR. Attention improves information flow between neuronal populations without changing the communication subspace. Curr Biol. 2021;31(23):5299–313 e4. doi: 10.1016/j.cub.2021.09.076.

22. MacDowell CJ, Buschman TJ. Low-Dimensional Spatiotemporal Dynamics Underlie Cortex-wide Neural Activity. Curr Biol. 2020;30(14):2665–80 e8. doi: 10.1016/j.cub.2020.04.090.

23. MacDowell CJ, Tafazoli S, Buschman TJ. A Goldilocks theory of cognitive control: Balancing precision and efficiency with low-dimensional control states. Curr Opin Neurobiol. 2022;76:102606. doi: 10.1016/j.conb.2022.102606.

24. Voytek B, Kayser AS, Badre D, Fegen D, Chang EF, Crone NE, et al. Oscillatory dynamics coordinating human frontal networks in support of goal maintenance. Nat Neurosci. 2015;18(9):1318–24. doi: 10.1038/nn.4071.

25. Brouwer GJ, Heeger DJ. Decoding and reconstructing color from responses in human visual cortex. J Neurosci. 2009;29(44):13992–4003. doi: 10.1523/JNEUROSCI.3577-09.2009.

26. Wolff MJ, Jochim J, Akyurek EG, Buschman TJ, Stokes MG. Drifting codes within a stable coding scheme for working memory. PLoS Biol. 2020;18(3):e3000625. doi: 10.1371/journal.pbio.3000625.

27. Panichello MF, Buschman TJ. Shared mechanisms underlie the control of working memory and attention. Nature. 2021;592(7855):601–5. doi: 10.1038/s41586-021-03390-w.

28. Xie Y, Hu P, Li J, Chen J, Song W, Wang XJ, et al. Geometry of sequence working memory in macaque prefrontal cortex. Science. 2022;375(6581):632–9. doi: 10.1126/science.abm0204.

29. Li HH, Curtis CE. Neural population dynamics of human working memory. Curr Biol. 2023. doi: 10.1016/j.cub.2023.07.067.

30. Kriegeskorte N, Wei XX. Neural tuning and representational geometry. Nat Rev Neurosci. 2021;22(11):703–18. doi: 10.1038/s41583-021-00502-3.

31. Ratcliffe O, Shapiro K, Staresina BP. Fronto-medial theta coordinates posterior maintenance of working memory content. Curr Biol. 2022;32(10):2121–9 e3. doi: 10.1016/j.cub.2022.03.045.

32. Glasser MF, Coalson TS, Robinson EC, Hacker CD, Harwell J, Yacoub E, et al. A multi-modal parcellation of human cerebral cortex. Nature. 2016;536(7615):171–8. doi: 10.1038/nature18933.

33. Smith SM. The future of FMRI connectivity. Neuroimage. 2012;62(2):1257–66. doi: 10.1016/j.neuroimage.2012.01.022.

34. Liegeois R, Santos A, Matta V, Van De Ville D, Sayed AH. Revisiting correlation-based functional connectivity and its relationship with structural connectivity. Netw Neurosci. 2020;4(4):1235–51. doi: 10.1162/netn_a_00166.

35. Rissman J, Gazzaley A, D’Esposito M. Measuring functional connectivity during distinct stages of a cognitive task. Neuroimage. 2004;23(2):752–63. doi: 10.1016/j.neuroimage.2004.06.035.

36. Conner AK, Briggs RG, Sali G, Rahimi M, Baker CM, Burks JD, et al. A Connectomic Atlas of the Human Cerebrum-Chapter 13: Tractographic Description of the Inferior Fronto-Occipital Fasciculus. Oper Neurosurg (Hagerstown). 2018;15(suppl_1):S436–S43. doi: 10.1093/ons/opy267.

37. Badre D, D’Esposito M. Functional magnetic resonance imaging evidence for a hierarchical organization of the prefrontal cortex. J Cogn Neurosci. 2007;19(12):2082–99. doi: 10.1162/jocn.2007.19.12.2082.

38. Badre D, Kayser AS, D’Esposito M. Frontal cortex and the discovery of abstract action rules. Neuron. 2010;66(2):315–26. doi: 10.1016/j.neuron.2010.03.025.

39. Cole MW, Ito T, Braver TS. The Behavioral Relevance of Task Information in Human Prefrontal Cortex. Cereb Cortex. 2016;26(6):2497–505. doi: 10.1093/cercor/bhv072.

40. Zhang J, Kriegeskorte N, Carlin JD, Rowe JB. Choosing the rules: distinct and overlapping frontoparietal representations of task rules for perceptual decisions. J Neurosci. 2013;33(29):11852–62. doi: 10.1523/JNEUROSCI.5193-12.2013.

41. Vaidya AR, Badre D. Abstract task representations for inference and control. Trends Cogn Sci. 2022;26(6):484–98. doi: 10.1016/j.tics.2022.03.009.

42. Schuck NW, Cai MB, Wilson RC, Niv Y. Human Orbitofrontal Cortex Represents a Cognitive Map of State Space. Neuron. 2016;91(6):1402–12. doi: 10.1016/j.neuron.2016.08.019.

43. Vaidya AR, Jones HM, Castillo J, Badre D. Neural representation of abstract task structure during generalization. Elife. 2021;10. doi: 10.7554/eLife.63226.

44. Zhou J, Jia C, Montesinos-Cartagena M, Gardner MPH, Zong W, Schoenbaum G. Evolving schema representations in orbitofrontal ensembles during learning. Nature. 2021;590(7847):606–11. doi: 10.1038/s41586-020-03061-2.

45. Zeithamova D, Dominick AL, Preston AR. Hippocampal and ventral medial prefrontal activation during retrieval-mediated learning supports novel inference. Neuron. 2012;75(1):168–79. doi: 10.1016/j.neuron.2012.05.010.

46. Morton NW, Schlichting ML, Preston AR. Representations of common event structure in medial temporal lobe and frontoparietal cortex support efficient inference. Proc Natl Acad Sci U S A. 2020;117(47):29338–45. doi: 10.1073/pnas.1912338117.

47. Fusi S, Miller EK, Rigotti M. Why neurons mix: high dimensionality for higher cognition. Curr Opin Neurobiol. 2016;37:66–74. doi: 10.1016/j.conb.2016.01.010.

48. Flesch T, Juechems K, Dumbalska T, Saxe A, Summerfield C. Orthogonal representations for robust context-dependent task performance in brains and neural networks. Neuron. 2022;110(7):1258–70 e11. doi: 10.1016/j.neuron.2022.01.005.

49. Kikumoto A, Mayr U. Conjunctive representations that integrate stimuli, responses, and rules are critical for action selection. Proc Natl Acad Sci U S A. 2020;117(19):10603–8. doi: 10.1073/pnas.1922166117.

50. Kikumoto A, Mayr U, Badre D. The role of conjunctive representations in prioritizing and selecting planned actions. Elife. 2022;11. doi: 10.7554/eLife.80153.

51. Gevins A, Smith ME, McEvoy L, Yu D. High-resolution EEG mapping of cortical activation related to working memory: effects of task difficulty, type of processing, and practice. Cereb Cortex. 1997;7(4):374–85. doi: 10.1093/cercor/7.4.374.

52. Hsieh LT, Ekstrom AD, Ranganath C. Neural oscillations associated with item and temporal order maintenance in working memory. J Neurosci. 2011;31(30):10803–10. doi: 10.1523/JNEUROSCI.0828-11.2011.

53. Hsieh LT, Ranganath C. Frontal midline theta oscillations during working memory maintenance and episodic encoding and retrieval. Neuroimage. 2014;85 Pt 2(02):721–9. doi: 10.1016/j.neuroimage.2013.08.003.

54. Liebe S, Hoerzer GM, Logothetis NK, Rainer G. Theta coupling between V4 and prefrontal cortex predicts visual short-term memory performance. Nat Neurosci. 2012;15(3):456–62, S1-2. doi: 10.1038/nn.3038.

55. Raghavachari S, Kahana MJ, Rizzuto DS, Caplan JB, Kirschen MP, Bourgeois B, et al. Gating of human theta oscillations by a working memory task. J Neurosci. 2001;21(9):3175–83. doi: 10.1523/JNEUROSCI.21-09-03175.2001.

56. Riddle J, Scimeca JM, Cellier D, Dhanani S, D’Esposito M. Causal Evidence for a Role of Theta and Alpha Oscillations in the Control of Working Memory. Curr Biol. 2020;30(9):1748–54 e4. doi: 10.1016/j.cub.2020.02.065.

57. Verbeke P, Ergo K, De Loof E, Verguts T. Learning to Synchronize: Midfrontal Theta Dynamics during Rule Switching. J Neurosci. 2021;41(7):1516–28. doi: 10.1523/JNEUROSCI.1874-20.2020.

58. Senoussi M, Verbeke P, Desender K, De Loof E, Talsma D, Verguts T. Theta oscillations shift towards optimal frequency for cognitive control. Nat Hum Behav. 2022;6(7):1000–13. doi: 10.1038/s41562-022-01335-5.

59. Wirsich J, Jorge J, Iannotti GR, Shamshiri EA, Grouiller F, Abreu R, et al. The relationship between EEG and fMRI connectomes is reproducible across simultaneous EEG-fMRI studies from 1.5T to 7T. Neuroimage. 2021;231:117864. doi: 10.1016/j.neuroimage.2021.117864.

60. Shi D, Yu Q. Distinct neural signatures underlying information manipulation in working memory. bioRxiv. 2023. doi: 10.1101/2023.07.11.548635.

61. Yu Q, Teng C, Postle BR. Different states of priority recruit different neural representations in visual working memory. PLoS Biol. 2020;18(6):e3000769. doi: 10.1371/journal.pbio.3000769.

62. Wan Q, Menendez JA, Postle BR. Priority-based transformations of stimulus representation in visual working memory. PLoS Comput Biol. 2022;18(6):e1009062. doi: 10.1371/journal.pcbi.1009062.

63. Cavanagh JF, Frank MJ. Frontal theta as a mechanism for cognitive control. Trends Cogn Sci. 2014;18(8):414–21. doi: 10.1016/j.tics.2014.04.012.

64. Konkle T, Brady TF, Alvarez GA, Oliva A. Conceptual distinctiveness supports detailed visual long-term memory for real-world objects. J Exp Psychol Gen. 2010;139(3):558–78. doi: 10.1037/a0019165.

65. Peirce J, Gray JR, Simpson S, MacAskill M, Hochenberger R, Sogo H, et al. PsychoPy2: Experiments in behavior made easy. Behav Res Methods. 2019;51(1):195–203. doi: 10.3758/s13428-018-01193-y.

66. Gramfort A, Luessi M, Larson E, Engemann DA, Strohmeier D, Brodbeck C, et al. MEG and EEG data analysis with MNE-Python. Front Neurosci. 2013;7:267. doi: 10.3389/fnins.2013.00267.

67. Esteban O, Markiewicz CJ, Blair RW, Moodie CA, Isik AI, Erramuzpe A, et al. fMRIPrep: a robust preprocessing pipeline for functional MRI. Nat Methods. 2019;16(1):111–6. doi: 10.1038/s41592-018-0235-4.

68. Gorgolewski K, Burns CD, Madison C, Clark D, Halchenko YO, Waskom ML, et al. Nipype: a flexible, lightweight and extensible neuroimaging data processing framework in python. Front Neuroinform. 2011;5:13. doi: 10.3389/fninf.2011.00013.

69. Li AY, Liang JC, Lee ACH, Barense MD. The validated circular shape space: Quantifying the visual similarity of shape. J Exp Psychol Gen. 2020;149(5):949–66. doi: 10.1037/xge0000693.

70. Kriegeskorte N, Mur M, Bandettini P. Representational similarity analysis - connecting the branches of systems neuroscience. Front Syst Neurosci. 2008;2:4. doi: 10.3389/neuro.06.004.2008.

71. Vinck M, Oostenveld R, van Wingerden M, Battaglia F, Pennartz CM. An improved index of phase-synchronization for electrophysiological data in the presence of volume-conduction, noise and sample-size bias. Neuroimage. 2011;55(4):1548–65. doi: 10.1016/j.neuroimage.2011.01.055.

